# Three-phase transitions to reproductive isolation: The roles of utilization mismatch and residual selection

**DOI:** 10.1101/595082

**Authors:** Géza Meszéna, Ulf Dieckmann

## Abstract

The adaptive emergence of reproductive isolation is increasingly recognized as a key mechanism of sympatric speciation. Here we aim at establishing a deeper understanding of the complex multilocus dynamics underlying such speciation transitions under resource competition. In reality, a single population’s resource utilization can never exactly match a resource distribution, making residual selection pressures inevitable. We find that this commonly leads to three-phase transitions to reproductive isolation. First, partial assortativity emerges, quickly adjusting a population’s variance to the resource distribution’s variance. Second, allelic variance slowly erodes across loci, allowing an increasingly bimodal phenotype distribution to emerge. Third, a fast transition occurs toward full bimodality in conjunction with practically complete reproductive isolation of the emerging two species. The first phase is driven by frequency-dependent divergent ecological selection. The second phase is driven by self-accelerating residual ecological selection: the more loci code for the selected phenotype, the slower is this intermediate phase. The third phase is driven by self-accelerating sexual selection. We study three types of mismatch-driven speciation, resulting from (i) incongruences between the shapes of resource distributions and competition kernels, (ii) low numbers of loci, and (iii) premature cessations of the first phase’s variance expansion. Our results suggest that the incomplete separation of incipient species, a characteristic of the second phase, is common in nature, which is likely resulting in detectable genetic footprints of three-phase transitions to reproductive isolation occurring in nature.

## Introduction

The emerging understanding of competitive, ecological, and adaptive speciation (Rosenzweig, 1978; Schluter, 2000; Dieckmann et al., 2004; Rundle and Nosil, 2005; Kopp et al., 2018) considers the origin of a new species as a transition in which ecological and genetic processes are linked in an intricate mutually dependent way. This understanding is empirically underpinned by the discovery of unequivocal cases of sympatric speciation (Schliewen et al, 1994; Via, 2001), as well as by ample field evidence for ecological adaptation and prolonged gene flow during speciation (Nosil, 2012). Speciation theory is challenged to describe and understand such complexity.

The selection regimes related to adaptation to an unexploited ecological opportunity are necessarily frequency-dependent, as fitness functions describing such situations inevitably depend on which niche is occupied and which is not (Rueffler et al., 2006). Adaptive dynamics theory (Metz et al., 1992, 1996; Dieckmann and Law, 1996; Geritz et al., 1998) provides a useful conceptual and mathematical framework for analyzing such situations, considering the consequences of frequency dependence in the absence of complications arising from sexual reproduction. Under frequency dependence, the adaptive evolution of a continuous character may cause it to converge to a fitness minimum (Eshel, 1983). The emerging divergent (i.e., frequency-dependent disruptive) selection pressures can lead to evolutionary branching, through which an asexual population splits into two subpopulations phenotypically evolving away from each other (Metz et al., 1992, 1996; Geritz et al., 1997, 1998; Meszéna et al., 1997, 2005; Doebeli and Dieckmann, 2000).

Adaptive-speciation models (e.g., Seger, 1985; Dieckmann and Doebeli, 1999; Kisdi and Geritz, 1999; Pennings et al., 2008) suggest that such disruptive selection may also select for the adaptive emergence of reproductive isolation in sexually reproducing organisms, eventually leading to speciation. While such models are considered convincing by some (e.g., Turelli et al., 2001), they have also met with significant criticism (e.g., Gavrilets, 2005). An important reason for skepticism has been the propensity of multilocus genetics to harbor large genetic variances, which may allow a population to exploit a wide resource distribution without speciation (Polechová and Barton, 2005; Bolnick and Fitzpatrick, 2007).

Our present paper’s analysis of the multilocus dynamics of adaptive speciation is centered on the simple but consequential observation that the match between an evolving phenotype distribution and the underlying resource distribution essentially is never perfect. The phenotype distribution of an unselected quantitative trait is Gaussian in the limit of infinitely many loci (infinitesimal model; Fischer, 1918, 1930; Bulmer, 1980; Turelli, 2017; Barton et al., 2017). Such a Gaussian phenotype distribution can be a perfect match for a resource distribution only under exceptional circumstances, such as in the Roughgarden model of character displacement, a Lotka-Volterra model often used to describe the ecology of adaptive speciation driven by resource competition based on the assumption of a Gaussian resource distribution and a Gaussian competition kernel (Roughgarden, 1979, pp. 534-536). In real life, however, the ingredient functions of such ecological models are not Gaussian and, independently, the infinitesimal limit is not necessarily relevant. Generically, we must therefore expect an imperfect match between resource production and resource consumption even after the evolutionary adjustment of a consumer’s population variance. The purpose of the current paper is to understand the wide-ranging consequences of such utilization mismatch. We investigate, in particular, how the residual selection emerging from a mismatch can contribute to a transition from a phenotypically wide and panmictic single population to a community comprising two or more phenotypically narrow and reproductively isolated populations.

To this end, we investigate the model by Dieckmann and Doebeli (1999) for significantly larger numbers of loci and higher population sizes. In this limit, we expect smoother and more deterministic behavior, more comparable to the well understood and intuition-shaping infinitesimal model. We study the ramifications of utilization mismatch primarily through non-Gaussian resource distributions and also explore other sources of mismatch that may affect speciation dynamics even when carrying capacities and competition kernels are both Gaussian. Given the high complexity of the resultant coupled ecological and genetic dynamics, we focus our analyses on assortativity determined by similarity in the focal ecological trait.

## Model

Following Dieckmann and Doebeli (1999), we consider diploid hermaphrodite individuals characterized by two quantitative traits, an ecological trait *x*_E_ and a mating trait *x*_M_ (the latter is called choosiness trait by Kopp et al., 2018). Both traits are determined by additive diallelic multilocus genetics, with *n*_E_ and *n*_M_ freely recombining diploid loci yielding 2*n*_E_ + 1 and 2*n*_M_ + 1 equidistant trait values, respectively. Allele reversal by mutation occurs with a small probability *μ* at reproduction. Population dynamics are governed by the per capita birth and death rates of individuals.

The allelic effects of the ecological trait are scaled such that its maximum allelic variance equals 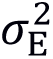 (Eq. 1a), while the allelic effects of the mating trait are scaled such that it ranges from −1 to +1 (Eq. 1b).

Individuals in their female role choose a mate and, if successful, produce a single offspring at rate *r*. Thus, the fertility of an individual in its female role depends on its probability of finding a mate, and the fertility of an individual in its male role depends on its propensity of being chosen as a mate. Mate choice depends on the individual’s mating trait *x*_M_, characterizing the type and degree of assortativity of mating with respect to the ecological trait: for assortatively mating individuals (*x*_M_ > 0), the probability of choosing a given individual as mating partner decreases with the difference in their ecological trait values (a “magic matching rule”; Servedio et al., 2011; Kopp et al., 2018), while the probability increases with this ecological difference for disassortatively mating individuals (*x*_M_ < 0), with the trait value *x*_M_ = 0 corresponding to random mating.

How ecological differences affect mating probabilities is described by a Gaussian function (Eq. 2a), whose standard deviation depends on *x*_M_ in a way controlled by a parameter *α* (Eq. 2b), as shown in Figure S1. We use *α* = 1/2, which implies that small trait values *x*_M_ > 0 cause a first-order reduction of mating probabilities (Eq. 2c; *α* = 2 was used by Dieckmann and Doebeli, 1999).

We assume that the probability of reproduction may be reduced by high choosiness, as it results in a scarcity of acceptable mates. This cost is measured by the parameter *c* (Eq. 2d), with *c* = 0 representing the absence of such a cost.

Individuals compete in an ecological setting described by Lotka-Volterra dynamics: the death rate of an individual with a given ecological phenotype is given by *r* times the ratio of the competition-effective population size and carrying capacity depending on that phenotype. The carrying-capacity distribution describes the latter dependence and represents the unloaded distribution of resources that are not explicitly modeled; it is therefore often referred to as the resource distribution. The competition-effective population size is the sum over all individuals, weighed by a competition kernel depending on the competing individuals’ ecological difference (Eq. 3a). Both the resource distribution and the competition kernel are symmetric unimodal functions characterized by standard deviations (*σ*_K_ and *σ*_C_, respectively) and kurtosis parameters (*k*_K_ and *k*_C_; Pigolotti et al., 2010; Leimar et al., 2013). Kurtosis parameters of 1 result in Gaussian functions, while kurtosis parameters larger than 1 result in platykurtic functions (Eq. 3b, Figure S2).

Model runs are initiated with all individuals having trait values *x*_E,i_ and *x*_M,i_. As the birth rate is set to *r* = 1, each individual on average produces one offspring per time unit, so the time unit equals the generation time. During a model run, we record the evolution of the population distributions of both traits, the allelic variance in each phenotype class of the ecological trait (Eq. 4a), and the distributions of ecological fitness (Eq. 5b) and male reproductive success (Eq. 5d) across the phenotype classes of the ecological trait, with the latter determining sexual fitness (Eq. 5c) and hence sexual selection. For each birth event, we record the difference between the ecological phenotypes of the parents, the Hamming distance between the corresponding ecological genotypes, as well as the deviation of the offspring’s ecological phenotype from the mid-parental phenotype.

The Appendix provides a full specification of the model, as well as of all quantities evaluated during model runs. Table 1 provides an overview of all model parameters.

**Table 1.** Model parameters.

| Description | Symbol | Equation |
| --- | --- | --- |
| Number of loci for the ecological trait | $n_E$ | 1a |
| Number of loci for the mating trait | $n_M$ | 1b |
| Maximum allelic standard deviation of the ecological trait | $\sigma_E$ | 1a |
| Per capita rate of choosing a mate | $r$ | |
| Parameters of the mating kernel | $s_+, s_-, \alpha$ | 2b |
| Cost of choosiness | $c$ | 2d |
| Standard deviation of the competition kernel | $\sigma_C$ | 3f |
| Kurtosis parameter of the competition kernel | $k_C$ | 3f |
| Maximum of the resource distribution | $K_0$ | 3g |
| Standard deviation of the resource distribution | $\sigma_K$ | 3g |
| Kurtosis parameter of the resource distribution | $k_K$ | 3g |
| Mutation probability | $\mu$ | |

## Results

We find that the multilocus dynamics underlying the adaptive emergence of reproductive isolation driven by frequency-dependent selection under resource competition commonly follow a characteristic pattern involving three phases. Below we show the robust occurrence of such three-phase transitions to reproductive isolation under a range of qualitatively different circumstances. In all cases, the mismatch-induced selection pressures that remain after a unimodal population distribution of the ecological trait has evolved as much as possible to match the resource distribution are key to understanding the transition patterns.

### Three-phase transitions for kurtosis-based mismatch

Figure 1 shows a three-phase transition to reproductive isolation for a slightly platykurtic resource distribution and a Gaussian competition kernel, with the boundaries between phases indicated by white vertical lines in the figure:

**Figure 1.**
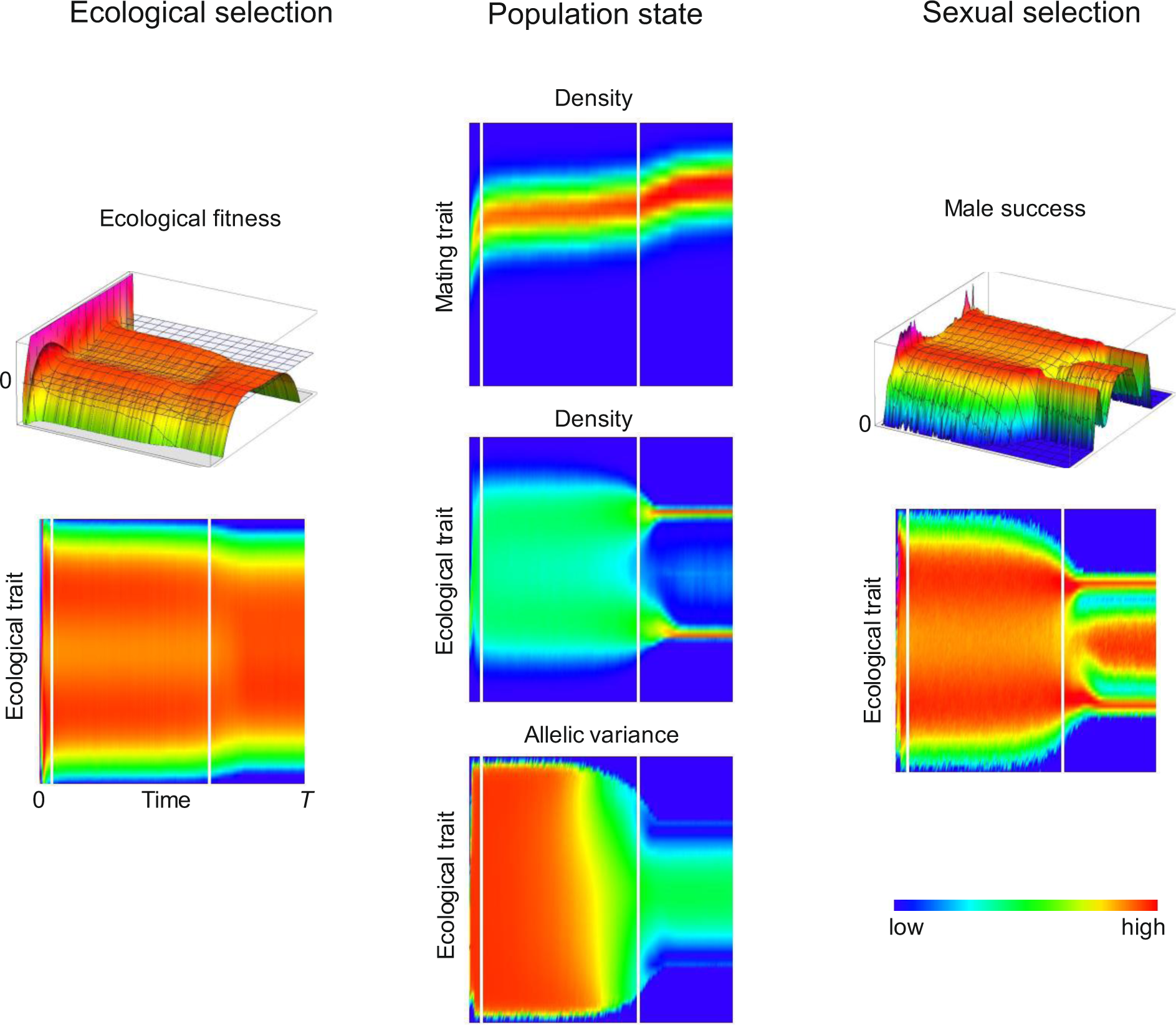
Three-phase transition to reproductive isolation. The central column of two-dimensional plots depicts the time course of the population state in terms of the population distribution of the mating trait (top), the population distribution of the ecological trait (middle), and the allelic variance of the ecological trait (bottom). The main determinants of selection pressures, i.e., the ecological fitness determining ecological selection (left column) and the male reproductive success determining sexual selection (right column), are shown both as two-dimensional plots and as three-dimensional landscapes. In all two-dimensional plots, the horizontal coordinate is time, while the vertical coordinate is either the ecological trait or the mating trait. Parameters: *n*_E_ = 32, *n*_M_ = 16, *σ*_E_ = 0.25, *r* = 1, *s*_+_ = 0.075, *s*_−_ = 0.5, *α* = 1/2, *c* = 0, *σ*_C_ = 1, *k*_C_ = 1, *K*_0_ = 10^5^, *σ*_K_ = 1, *k*_K_ = 1.6, *μ* = 10^−4^, *x*_E,i_ = −0.85, *x*_M,i_ = 0, and *T* = 2,000.

### First phase

The population mean of the ecological trait evolves to where the resource distribution is maximal, after which the population mean of the mating trait increases to a positive value corresponding to assortative mating, allowing the population variance of the ecological trait to increase. Throughout this phase, the population distributions of both traits remain unimodal.

### Second phase

Genetic diversity, measured by the allelic variances of the ecological trait within each of its phenotype classes, gradually diminishes in a slowly starting yet self-accelerating process. In accordance with these dynamics, the population distribution of the ecological trait gradually turns bimodal and the population mean of the mating trait increases further.

### Third phase

Once genetic diversity falls below a critical level, the ecological trait undergoes a fast transition to full bimodality and the population mean of the mating trait increases further. At the end of this phase, the rate of hybridization between the two narrow modes in the population distribution of the ecological trait is so low that these are practically completely reproductively isolated, indicating speciation.

Figure S3 provides additional information about key quantities involved in the three-phase transition, and Figure 5a shows the ecological fitness at the end of the first phase, quantifying the aforementioned mismatch between the resource distribution and the distribution of competition-effective population sizes resulting for a unimodal population distribution of the ecological trait. We will return to this information when offering in the Discussion a detailed process-based interpretation of the three-phase transition.

### Robustness of three-phase transitions

To investigate the robustness of the observed three-phase transition, Figures 2, 3, and S4 show the effects of varying model parameters relative to their reference values used in Figure 1.

**Figure 2.**
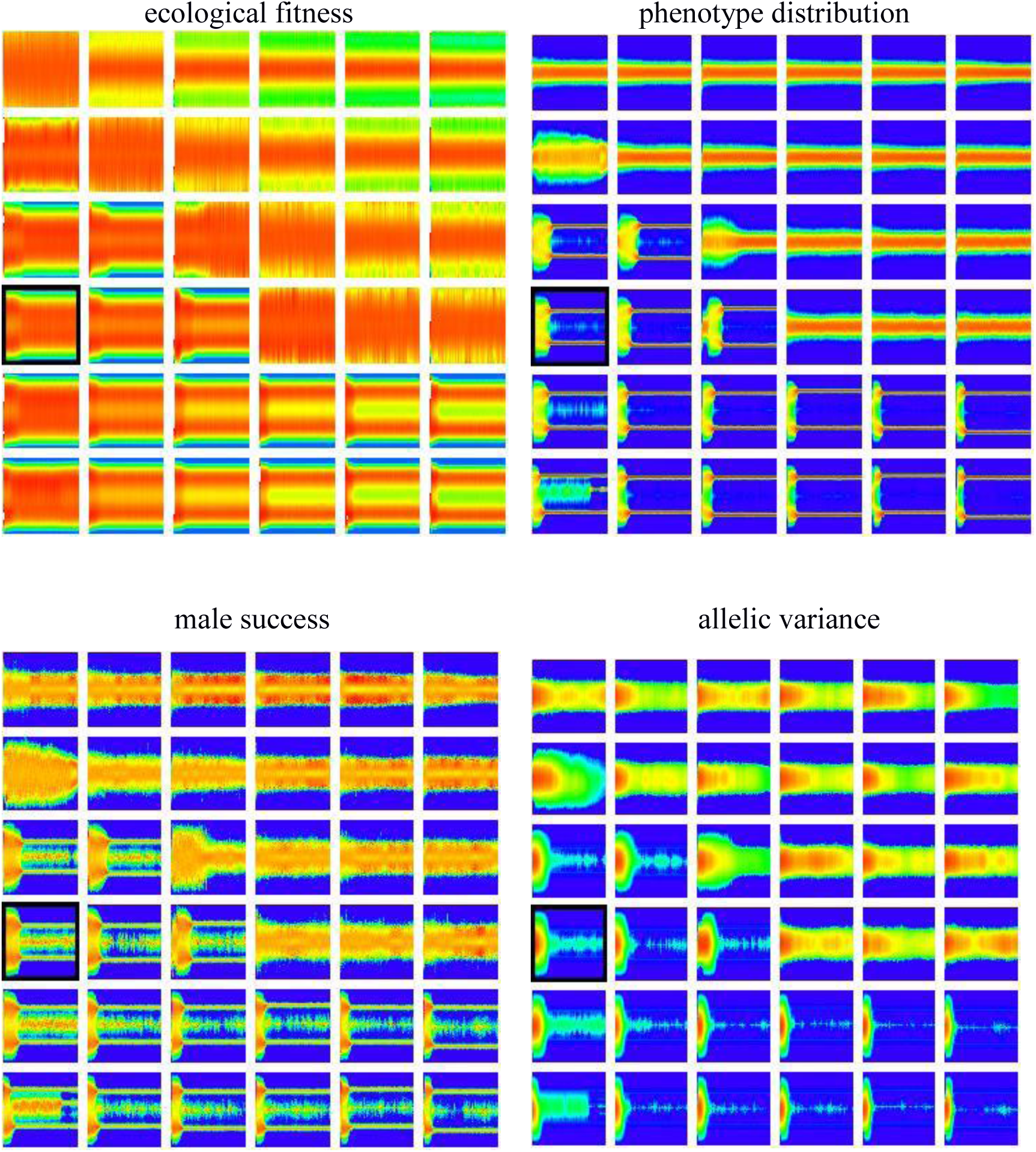
Dependence of three-phase transitions to reproductive isolation on the kurtosis parameter *k*_K_ of the resource distribution and the kurtosis parameter *k*_C_ of the competition kernel. In each panel, the horizontal coordinate is time, while the vertical coordinate is the ecological trait. In the four matrices of panels, the color coding indicates the ecological fitness (top left), the population distribution of the ecological trait (top right), the male reproductive success (bottom left), or the allelic variance of the ecological trait (bottom right). In each of the four matrices of panels, *k*_K_ = 1, 1.2, 1.4, 1.6, 1.8, 2 increases in the rows of panels from top to bottom, while *k*_C_ = 1, 1.2, 1.4, 1.6, 1.8, 2 increases in the columns of panels from left to right. Other parameters are as in Figure 1, except for *K*_0_ = 10^4^ and *T* = 5,000. The four panels framed in black indicate the combination of kurtosis parameters used in Figure 1.

**Figure 3.**
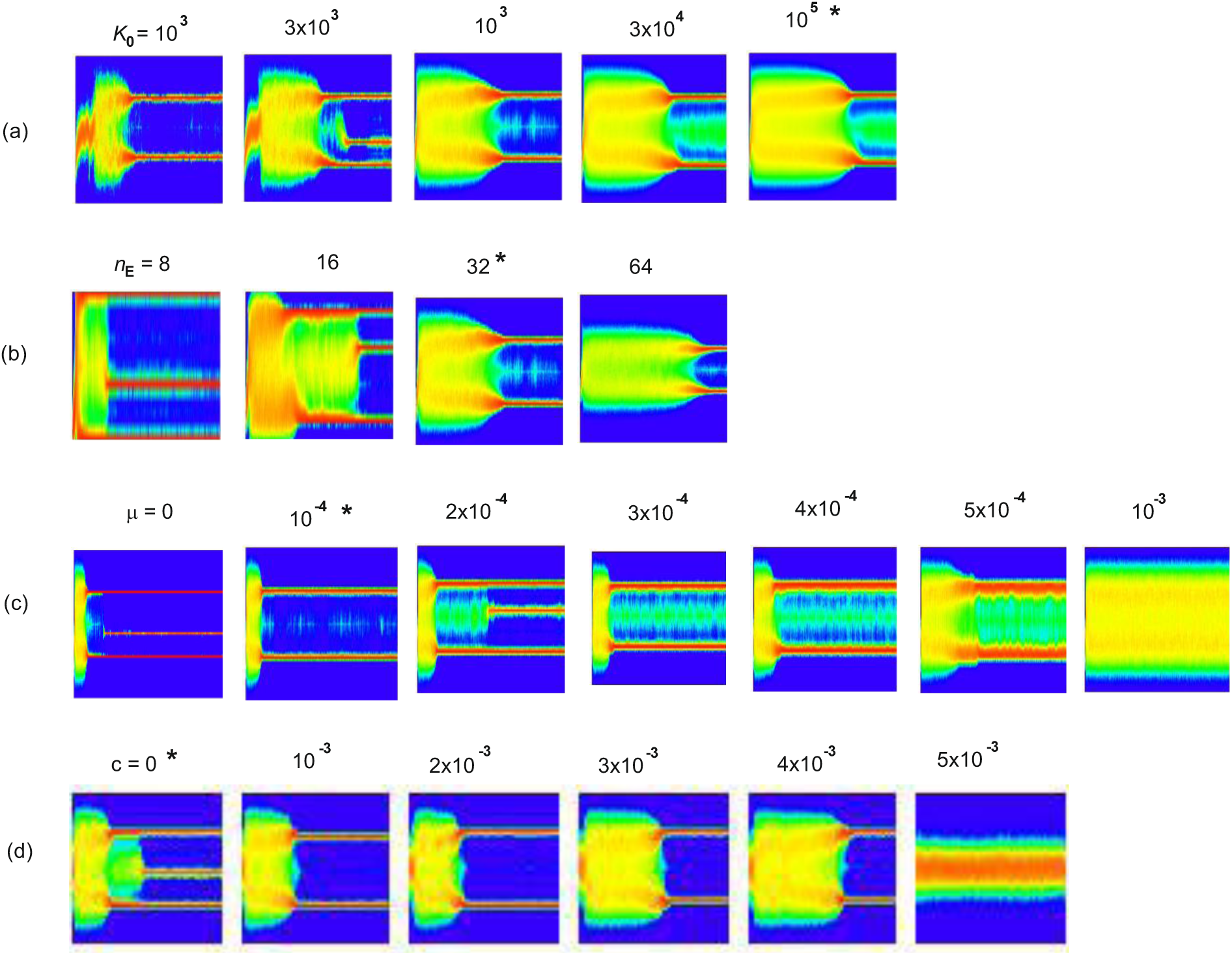
Dependence of three-phase transitions to reproductive isolation on (a) population size as scaled by the maximum carrying capacity *K*_0_, (b) the number *n*_E_ of ecological loci, (c) the mutation probability *μ*, and (d) the cost *c* of choosiness. In each panel, the horizontal coordinate is time, while the vertical coordinate is the ecological trait. Other parameters are as in Figure 1, except for *K*_0_ = 10^4^ and (c, d) *T* = 5,000. The four asterisks indicate the parameter values used in Figure 1.

Figure 2 illustrates how the kurtoses of the resource distribution and of the competition kernel influence the occurrence and dynamics of three-phase transitions. Making the resource distribution more platykurtic, by increasing the kurtosis parameter *k*_K_, facilitates speciation and makes it occur faster, while changing the kurtosis parameter *k*_C_ of the competition kernel has the opposite effect. The inequality *k*_K_ > *k*_C_ serves as a rough criterion for speciation to occur, in line with expectations. After the population variance in the ecological trait has temporarily equilibrated at the end of the first phase, an excess platykurtosis of the resource distribution relative to the competition kernel provides excess resources at the edges of the population distribution of the ecological trait, favoring extreme phenotypes of the ecological trait at the end of the first phase. An excess platykurtosis of the competition kernel has the opposite effect, favoring intermediate phenotypes of the ecological trait at the end of the first phase. These observations underscore the importance of the selection pressures remaining after a unimodal population distribution of the ecological trait has evolved as much as possible to match the resource distribution, in terms of the resultant distribution of competition-effective population sizes. As we will see in more detail below, appreciating the role of such utilization mismatch, and of the residual selection it entails, is central to understanding the evolutionary mechanisms underlying three-phase transitions to reproductive isolation.

Figure 3 examines the effects of population size, which is scaled by *K*_0_, of the number *n*_E_ of loci for the ecological trait, of the mutation probability *μ*, and of the cost *c* of choosiness. We find that the duration of the second phase is only mildly affected by population size: even when population size is raised by two orders of magnitude, this duration increases only modestly, by a factor of 2-3 (Figure 3a). Finding this weak dependence is very significant, as it shows that the loss of genetic diversity during the second phase is driven by selection, and not merely by random genetic drift. In contrast, there is a clear dependence of the second phase’s duration on the number of loci for the ecological trait: more loci imply a longer second phase and smoother, more regular dynamics (Figure 3b). This observation will be essential for understanding the slowness of the second phase in the Discussion. Increasing the mutation probability (Figure 3c) or the cost of chosiness (Figure 3d) slows down the second phase. In both cases, the progress toward speciation becomes fully arrested beyond threshold values of these parameters.

Figure S4a depicts the dependences on the standard deviation *σ*_C_ of the competition kernel and on the maximum allelic standard deviation *σ*_E_ of the ecological trait. We find that speciation does not occur, or occurs with a delay, for extremely small *σ*_E_. When *σ*_E_ is large enough for speciation to occur, its further increase makes the second phase longer, while very large values of *σ*_E_ prevent speciation. Decreasing *σ*_C_ enables the emergence of more than two species, either simultaneously, or via hybridization of two already existing, but not completely reproductively isolated, species (Bolnick, 2006). Very small values of *σ*_C_, instead of leading to very many similar yet reproductively isolated species, prevent speciation. Figure S4b depicts the aforementioned dependences for *α* = 2, in which case speciation happens for a narrower parameter range and in a different way, as discussed in the next subsection.

We conclude that, while details of the three-phase transitions depend on many parameters in complicated ways, both the final outcome of practically complete reproductive isolation among two or more species and the characteristic three phases of these transitions robustly occur across wide parameter ranges. The observed dependencies suggest that the rate-limiting step in the three-phase transitions is the second phase, during which residual selection drives the gradual elimination of genetic diversity.

### Three-phase transitions for the doubly Gaussian case

The dependence on the kurtosis parameters shown in Figure 2 could be misinterpreted as suggesting an absence of speciation dynamics when the resource distribution and the competition kernel are both Gaussian, *k*_K_ = *k*_C_ = 1, which we refer to as the doubly Gaussian case. In this case, it would be possible, in principle, for the mismatch between resource production and resource consumption to disappear at a particular value of the population variance in the ecological trait. The reason is that resource consumption in the Roughgarden model is described by the distribution of competition-effective population sizes, i.e., by the convolution of the population distribution of the ecological trait with the competition kernel (Eq. 3a). If both functions are Gaussian, their convolution is also Gaussian, which could, in principle, result in a full match with a Gaussian resource distribution. If this happened, it would imply, already at the end of the first phase, a cessation of all selection pressures resulting from resource competition.

Interestingly, Figures 4a and S5 present clear speciation dynamics in a specific model run for a doubly Gaussian case with a smaller *σ*_E_ than in Figure 1. As before, population sizes and numbers of loci are sufficiently large to enable smooth and mostly deterministic dynamics. For the shown parameter combination, the evolution of increased assortativity and the accompanied increase of population variance in the ecological trait do not commence. This causes a utilization mismatch favoring extreme phenotypes of the ecological trait, as shown in Figure 5b. The resultant unimodal population distribution with high mismatch is metastable: after a certain waiting time, a sudden shift occurs in the population distribution of the mating trait, leading to a suddenly increased population variance in the ecological trait, which ends the first phase. This is followed by the second and third phase as before, which completes the transition to reproductive isolation.

**Figure 4.**
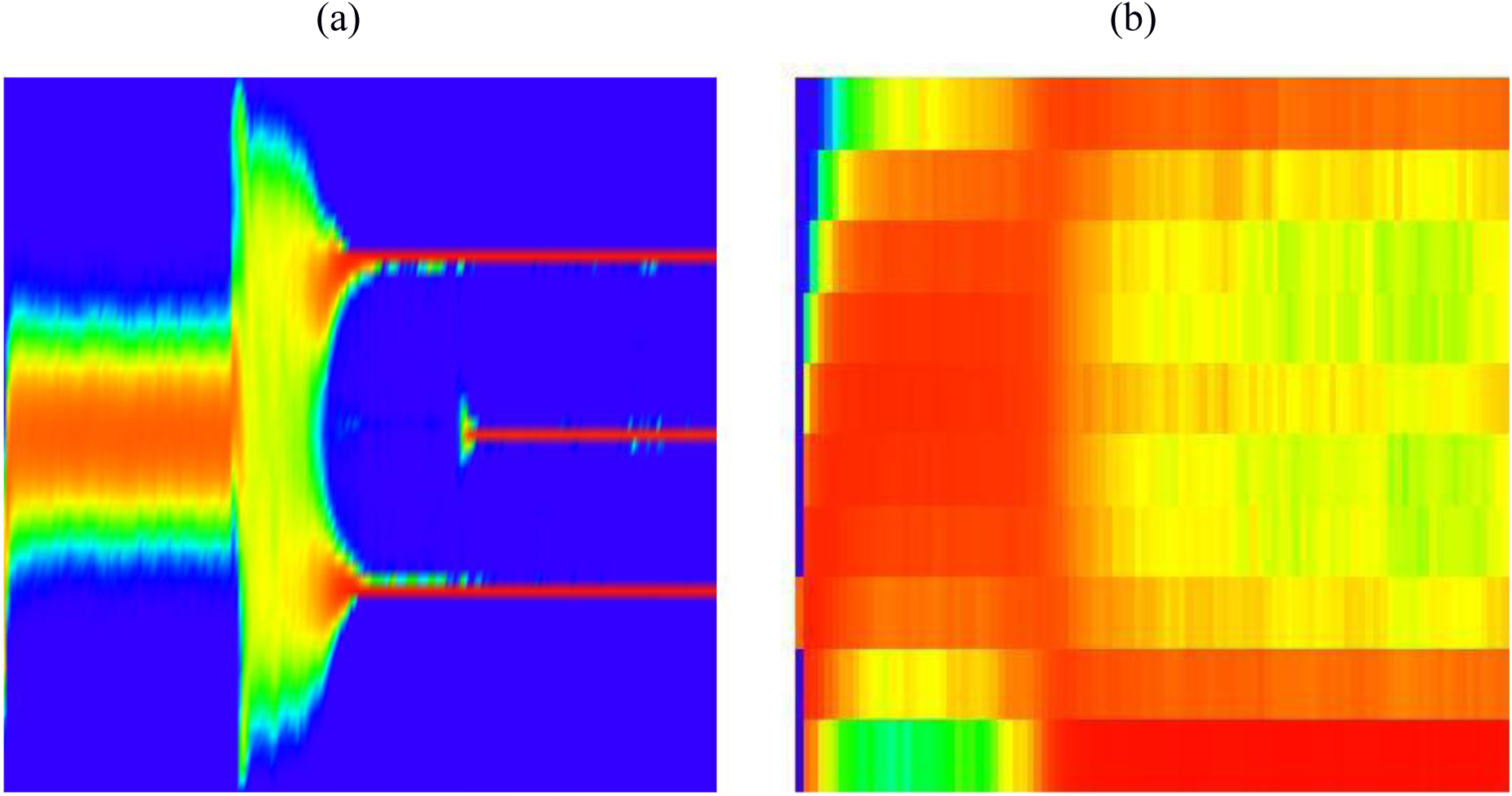
Three-phase transition to reproductive isolation in the doubly Gaussian case, *k*_K_ = *k*_C_ = 1. In both panels, the horizontal coordinate is time, while the vertical coordinate is the ecological trait. (a) Utilization mismatch resulting from a low maximum allelic variance 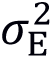 of the ecological trait, causing fluctuation-induced speciation dynamics for large population sizes and high numbers of loci. (b) Utilization mismatch resulting from a low number *n*E of loci for the ecological trait, causing speciation dynamics leading to population distributions of the ecological trait dominated by the extreme phenotypes. Other parameters are as in Figure 1, except for *K*_0_ = 10^4^, *σ*_A_ = 0.158, *σ*_C_ = 0.75, and (a) *μ* = 0, *T* = 10,000, (b) *n*_E_ = *n*_M_ = 5, *s*_+_ = 0.05, *s*_-_ = 1, *α* = 2, *μ* = 10^−3^, *T* = 500.

Figure S6 depicts replicate model runs for the same parameter combination, with individual runs differing only in their random seed. We find that, for the doubly Gaussian case with large population sizes and large numbers of loci, the waiting time for the end of the first phase is probabilistically distributed, which is consistent with it being initiated by a sufficiently large fluctuation. In some model runs, waiting for this fluctuation-induced transition to high population variance in the ecological trait causes the population to lose its genetic diversity while remaining unimodal. Consequently, the window for fluctuation-induced speciation dynamics in the doubly Gaussian case is not open indefinitely.

The described fluctuation-induced speciation dynamics are not restricted to the doubly Gaussian case. Figure S4b shows replicate model runs for a platykurtic resource distribution that include multiple examples that, together, show how the first phase ends after a probabilistically distributed waiting time. In general, fluctuation-induced speciation dynamics may occur when the initial increase of assortativity is arrested.

### Three-phase transitions for small numbers of loci

Figure 4b presents another variant of three-phase transitions for the doubly Gaussian case, which occur when the number of loci for the ecological trait is small. Under these circumstances, the population distribution of the ecological trait cannot be close to Gaussian, as it involves only a small number of phenotype classes and is restricted to a finite phenotype interval. Figure 5c shows the resulting mismatch: the ecological fitness is maximal at the extreme phenotypes even when the population distribution of the ecological trait is wide. The parametrization of this model run corresponds to those in Dieckmann and Doebeli (1999), except that here we use a significantly higher population size.

**Figure 5.**
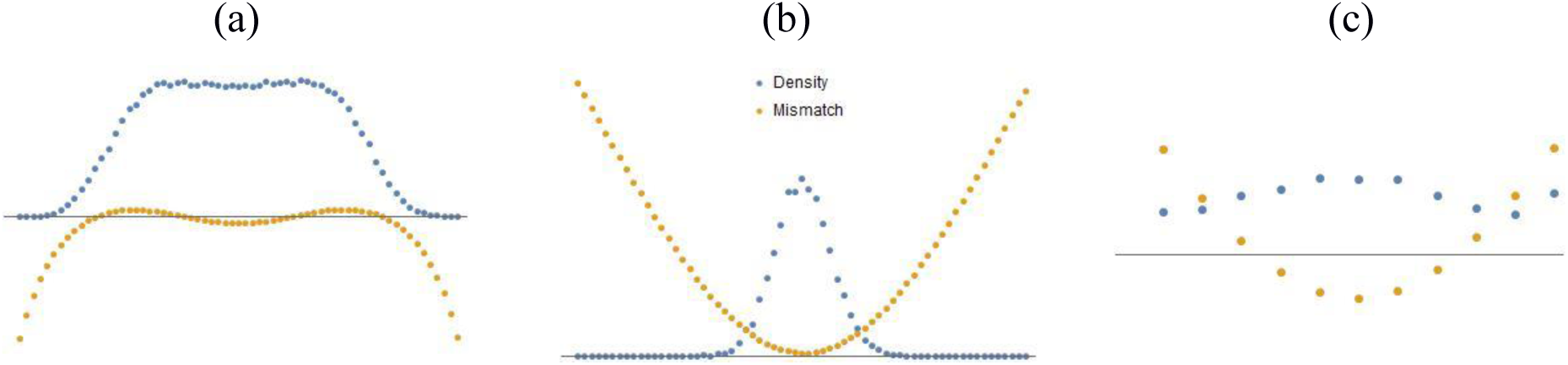
Utilization mismatches originating from diverse sources, all driving three-phase transitions to reproductive isolation. In each panel, the horizontal axis shows the ecological trait, while the vertical axes show the population distribution of the ecological trait (blue) and the corresponding ecological fitness (brown) at the end of the first phase (denoted by *T*\*). (a) Mismatch resulting from a resource distribution that is more platykurtic than the competition kernel, *k*_K_ > *k*_C_ (Figure 1, *T*\* = 100). (b) Mismatch in the doubly Gaussian case resulting from a low maximum allelic variance 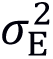 of the ecological trait (Figure 4a, *T*\* = 3,000). (c) Mismatch in the doubly Gaussian case resulting from a low number *n*_E_ of loci for the ecological trait (Figure 4b, *T*\* = 25). Parameters are as in Figures 1, 4b, and 4c, respectively.

In Figure S7, we examine how this type of three-phase transition depends on model parameters. Characteristically, the emerging bimodal population distributions of the ecological trait always involve a dominance of the two extreme phenotypes. For small population sizes, the dynamics are highly stochastic (occasionally even exhibiting back-and-forth transitions between unimodal and multimodal population distributions of the ecological trait). For large population sizes, speciation is contingent on small numbers of loci. For small numbers of loci, speciation occurs robustly, even when population sizes are large.

## Discussion

Our study seeks to contribute to a deeper understanding of adaptive speciation driven by frequency-dependent selection. Why does this process lead to different species, as opposed to a single species with high genetic variance (Polechová and Barton, 2005; Bolnick and Fitzpatrick, 2007)? To address this question, we have investigated the model by Dieckmann and Doebeli (1999) for significantly larger numbers of loci and higher population sizes.

Our main conclusion is that residual selection pressures, which under resource competition readily arise from a mismatch between resource production and resource consumption, are key for driving the speciation process and often cause a characteristic three-phase transition to reproductive isolation. Under such conditions, the speciation process is initiated by the expansion of population variance in the ecological trait governing resource competition, enabled by a first evolutionary increase of assortative mating (first phase). This is followed by a slowly growing deformation of the population distribution of the ecological trait from a unimodal shape to a bimodal shape, enabled by the initially slow but self-accelerating selection-driven elimination of allelic variance of the ecological trait and a second evolutionary increase of assortative mating (second phase). Speciation is concluded by a sharp transition to practically complete reproductive isolation of two ecologically differentiated populations, enabled by self-accelerating sexual selection and a third evolutionary increase of assortative mating (third phase). We have shown that the utilization mismatches that lie at the heart of this process can arise from different sources: incongruences between the shapes of resource distributions and competition kernels, low numbers of loci, and premature cessations of the first phase’s variance expansion.

In retrospect, the phased progress toward speciation was discernible already in an earlier study by Doebeli and Dieckmann (2003, their Figure 2b). A two-step process is clearly present also in the two-locus dynamics investigated by Rettelbach et al. (2011). The higher resolution and more deterministic behavior of the model analyzed here, permitted by studying larger numbers of loci and higher population sizes, critically help to reveal the three phases of the speciation process arising under these conditions.

Below we discuss the inferred mechanisms underlying three-phase transitions, the wider theoretical context, and the implications for understanding speciation. We do so based on the conceptual framework depicted in Figure 6. The population state is represented by three descriptors: the population distribution of the ecological trait, the allelic variance of this trait across its phenotype classes, and the population distribution of the mating trait. The first two descriptors are roughly independent, as analytically shown for large numbers of loci (Barton et al., 2017). The last two descriptors are roughly homogeneous (in the sense that allelic variances of the ecological trait are similar across its phenotype classes and that the population distribution of the mating trait is narrow), as numerically shown by our analyses. The selection pressures operating on the population state result from a frequency-dependent fitness landscape determined additively by fitness components describing ecological selection and sexual selection, with the latter component determined by male reproductive success. For the sake of brevity, we refer to these two components as ecological fitness and sexual fitness, respectively.

**Figure 6.**
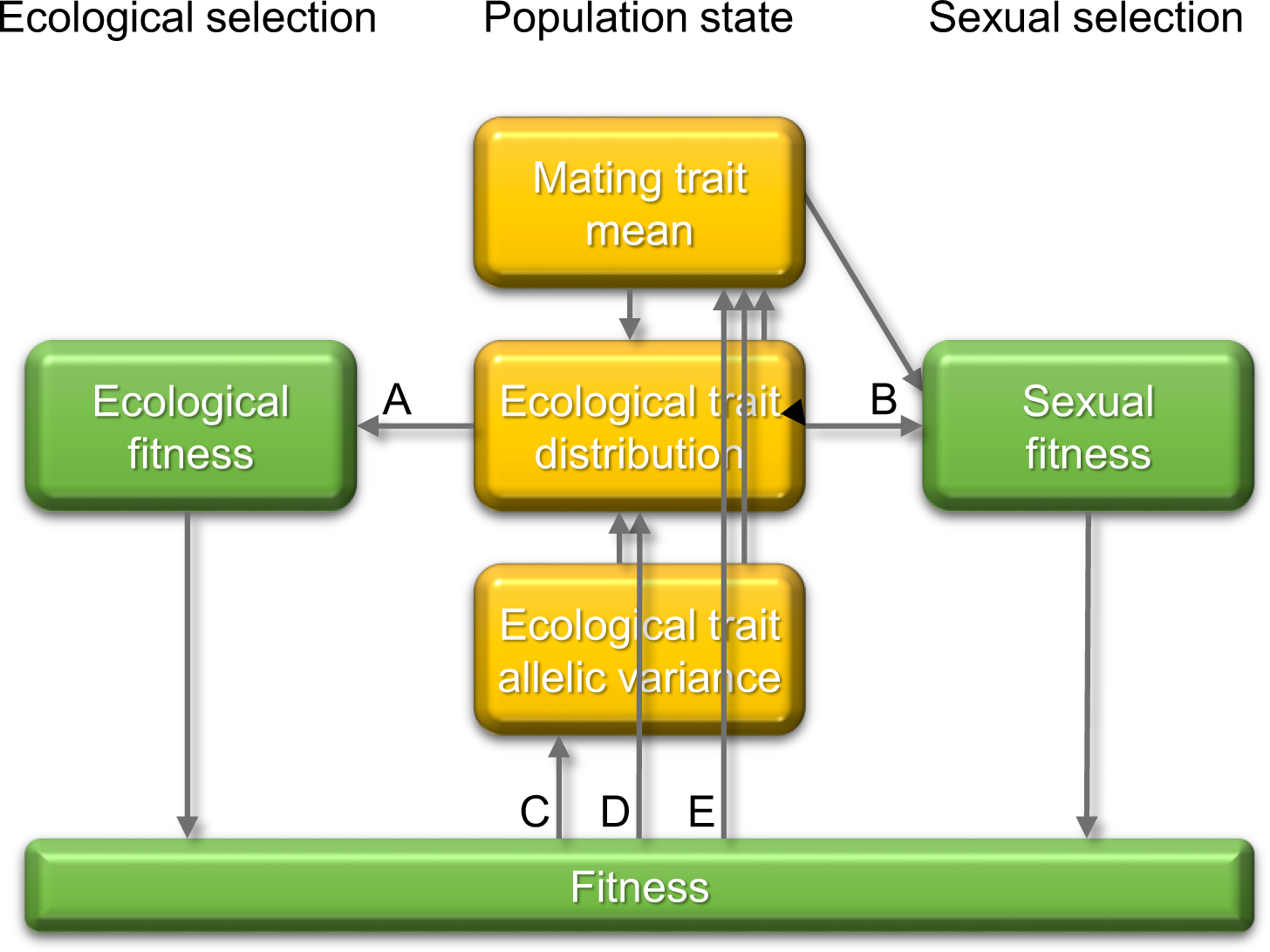
Conceptual framework for understanding the mechanisms underlying three-phase transitions to reproductive isolation. Yellow boxes show the salient descriptors of the population state: the population mean of the mating trait (top), the population distribution of the ecological trait (middle), and the average allelic variance of that trait across its phenotype classes (bottom). Green boxes show the components of fitness, determining ecological selection (left) and sexual selection (right). Main causal effects are indicated by arrows. A: Ecological fitness is determined by the population distribution of the ecological trait. B: Sexual fitness (through male reproductive success) is jointly determined by the population distribution of the ecological trait and the population mean of the mating trait. C: Evolution of the average allelic variance of the ecological trait is determined by the average curvature of the fitness landscape. D: Evolution of the population distribution of the ecological trait is jointly determined by the fitness landscape, the population mean of the mating trait, and the average allelic variance of the ecological trait. E: Evolution of the population mean of the mating trait is jointly determined by the fitness landscape, the population distribution of the ecological trait, and the average allelic variance of the ecological trait.

### Ecological selection drives processes traditionally associated with stabilizing and disruptive selection

Selection against intermediate phenotypes of the ecological trait is expected to select for ecological bimodality (Effect D in Figure 6) and assortative mating (Effect E in Figure 6), and thus for reproductive isolation (see, e.g., Pennings et al., 2008, for a minimal, analytically tractable model). However, there is a constraint on this process: assortative mating is based on ecological phenotypes, instead of the genotypes of multilocus quantitative characters. Therefore, assortative mating alone cannot lead to reproductive isolation while segregation variance remains high. Segregation variance measures the variance of offspring phenotypes around mid-parental phenotypes and thus naturally increases with allelic variance. In this situation, evolution toward reproductive isolation must be a combined process, in which assortativity increases and allelic variance decreases. We propose that the slow selection-driven loss of allelic variance of the ecological trait is the rate-limiting step of this combined process: the larger the number of loci for the ecological trait, the longer is this step, and hence the second phase of the three-phase transition (Figure 3ab). In Haken’s (1983) terminology, changes in the population mean of the mating trait and in the shape of the population distribution of the ecological trait are enslaved by the slow process of selection-driven erosion of allelic variance in the ecological trait.

Neither of the standard approximations of multilocus genetic – the infinitesimal model (Fisher, 1918; Turelli, 2017; Barton et al., 2017) and the hypergeometric model (also known as the symmetric model; Barton, 1992; Doebeli, 1996a, 1996b; Kondrashov and Kondrashov, 1999; Sachdeva and Barton, 2017) – describe the selection-driven loss of genetic variance. In the first approximation, allelic variance is not affected by selection. In the second approximation, equal allele frequencies are assumed across all loci (implying maximal allelic variance given the population mean). For establishing a detailed process-based understanding of the three-phase transitions, we use the infinitesimal model (i.e., the limit of infinitely many loci) as a reference point.

For large, but not infinite, numbers of loci, the selection-driven change of a trait’s allelic variance is determined by the average curvature (second derivative) of the corresponding phenotype’s fitness landscape (Bulmer, 1980, p. 166), where the average has to be taken according to the phenotype’s population distribution. In particular, allelic variance decreases when this average curvature is negative, and increases when it is positive. Moreover, the rate of selection-driven change in allelic variance is inversely proportional to the number of loci: consequently, this change becomes slow when the number of loci is large, and vanishes in the infinitesimal limit.

In our numerical analyses, we have observed a roughly constant negative average curvature of ecological fitness. According to the aforementioned theoretical results, this is expected to cause the selection-driven elimination of allelic variance during the second phase of the three-phase transitions (Effect C in Figure 6), in agreement with our observations (Figure S3a). The negative sign of the average curvature is understandable, because averaging according to the bimodal population distribution of the ecological trait emphasizes the parts of the fitness landscape that are surrounding its two modes (uniform averaging would lead to zero average curvature). Two other observations – the lengthening of the second phase for larger numbers of loci (Figure 3b) and for higher maximum allelic variances of the ecological trait (Figure S4a) – are also in line with the theoretical expectation. The selection-driven loss of allelic variance is a departure from the unstable equilibrium of equal allele frequencies, where the hypergeometric model is applicable. Therefore, this variance loss starts slowly and initially self-accelerates exponentially with growing distance from the unstable equilibrium. Eventually, allelic variance converges to zero, which is the stable equilibrium point of the dynamics of allelic variance under negative average curvature (Figure S3b), again in accordance with the theoretical expectation.

The joint occurrence of selection for ecological bimodality (Effect D in Figure 6) and assortative mating (Effect E in Figure 6) on the one hand with selection for reduced allelic variance (Effect C in Figure 6) on the other hand is surprising, because they are traditionally associated with disruptive and stabilizing selection, respectively. However, the distinction between disruptive and stabilizing selection is unequivocal only for quadratic fitness landscapes. In contrast, the bimodal fitness landscapes emerging in the speciation processes studied here are far from quadratic and can thus drive processes simultaneously that for quadratic fitness landscapes can occur only separately. On the one hand, they have a disruptive nature, selecting against intermediate phenotypes of the ecological trait and thus for ecological bimodality and assortative mating. On the other hand, they also have a stabilizing nature, selecting for reduced allelic variance. Importantly, this mixed selection regime, which is key to the emergence of reproductive isolation through three-phase transitions, arises naturally from the underlying ecology of resource competition.

### Sexual selection drives runaway evolution

Sexual selection is an unavoidable consequence of assortative mating. With female mate choice, an individual’s reproductive success in its male role is proportional to the number of individuals in their female role ready to choose it as a mate. With assortative mating, the ecological phenotypes that are most abundant, together with phenotypes similar to them, thus enjoy a reproductive advantage. This advantage of commonality leads to the “instability of the sexual continuum”, which was analytically studied by Noest (1997) in a model with fixed segregation variance, corresponding to the infinitesimal limit, and a fixed degree of assortative mating.

In our model, Noest’s instability is augmented by two positive feedback loops arising from evolving allelic variance, and thus evolving segregation variance. The first feedback loop operates through the population distribution of the ecological trait (BC in Figure 6). As the more common phenotypes of the ecological trait enjoy an advantage from sexual selection, sexual fitness has a negative average curvature, and therefore always selects for reduced allelic variance of the ecological trait (Effect C in Figure 6). This, in turn, narrows the modes of the population distribution of the ecological trait, which makes the variance-decreasing sexual selection even stronger (Effect B in Figure 6).

The second feedback loop operates through the evolving mating trait (BE in Figure 6). A decreasing allelic variance of the ecological trait shifts the optimal population mean of the mating trait to higher values (Effect E in Figure 6), implying stronger assortativity, which makes the variance-decreasing sexual selection even stronger (Effect B in Figure 6).

Through these two feedbacks, sexual selection results in runaway evolution. This prediction is corroborated by our numerical analyses: while the negative average curvature of ecological fitness increases in absolute value only slowly during the second phase of the speciation process, the negative average curvature of sexual fitness becomes larger and larger in absolute value throughout this phase (Figure S3a).

We highlight that the described runaway evolution is different from the well-known Fisherian runaway (Fisher 1930), which results from a positive feedback loop between a male trait evolving jointly with a female preference for it. We also mention the study by Doebeli et al. (2007), which extended the work by Noest (1997) to non-uniform resource distributions, non-Gaussian competition kernels, and evolving degrees of assortative mating, while treating segregation variance as a model parameter, rather than as being subject to evolution, as in our present analyses.

### Detailed process-based understanding of the three-phase transition pattern

We now have all ingredients in place to offer a detailed process-based understanding of the three-phase speciation process as illustrated in Figure 1. As explained above, the three salient descriptors of the population state are the population distribution of the ecological trait, the allelic variance of this trait, and the population mean of the mating trait, with selection governed by a fitness landscape given by the sum of ecological fitness and sexual fitness.

The *first phase* of the speciation process can be understood as the initial relaxation of the population’s trait distributions on a fast timescale at constant allelic variance. The process starts with the directional evolution of the population mean of the ecological trait to the peak of the resource distribution. At this point, the ecological fitness is bimodal owing to the effects of frequency-dependent competition. The associated ecological selection against intermediate phenotypes of the ecological trait results in directional selection on the mating character for higher assortativity (Effect E in Figure 6). In turn, higher assortativity results in increased population variance of the ecological trait, while its population distribution remains roughly Gaussian. This process is fast, because it involves only directional selection. It stops when the population variance of the ecological trait becomes wide enough for resource consumption to match resource production as well as possible: at this point, a further rise of assortativity, and thus of population variance in the ecological trait, would strengthen selection against the extreme phenotypes of the ecological trait, resulting in selection for decreased assortativity. Still, because of the platykurtosis of the resource distribution, a mismatch remains between resource consumption and resource production, resulting in residual selection pressures; i.e., ecological selection does not cease at the end of the first phase.

The *second phase* is characterized by the slow, but self-accelerating, selection-driven loss of allelic variance, caused by the negative average curvature of both ecological fitness and sexual fitness (Effect C in Figure 6). The enslaved population distribution of the ecological trait follows this change (Effect D in Figure 6), subject to two opposing forces: selection pushes it toward bimodality, against the smoothing effects of hybridization, segregation, and recombination. Likewise, the enslaved population mean of the mating trait follows the loss of allelic variance (Effect E in Figure 6). With decreasing allelic variance, the population becomes more and more bimodal and concentrated around the fitness peaks.

The details of this process are shown in Figure S3b. Decreasing allelic variance decreases the average Hamming distance between parental genotypes (Figure S3b), which in turn decreases the segregation variance of offspring phenotypes around mid-parental phenotypes (Figure S3b), which in turn increases the advantage (avoidance of intermediate offspring phenotypes) and decreases the disadvantage (production of extreme offspring phenotypes) of phenotype-based assortative mating, which in turn increases the population mean of the mating trait (Figure 1), which in turn decreases the average squared difference of the ecological trait in mating pairs (Figure S3b), which in turn contributes to the emergence of bimodality in the population distribution of the ecological trait (Figure 1). The average Hamming distance and segregation variance are proportional to each other and follow a time course similar to that of the allelic variance. Interestingly, the time courses of the genetic differences between parents (measured by the average Hamming distance in mating pairs) and the phenotypic differences between them (measured by the average squared difference of the ecological trait in mating pairs) are very different: this is in line with the theoretical expectation that the allele frequency at any single locus becomes independent from the corresponding phenotype when the number of loci becomes large (Barton et al., 2017).

The *third phase* is markedly different and is initiated when the self-accelerating sexual selection starts to dominate (Figure S3a). The two associated positive feedback loops described above (BC and BE in Figure 6) cause a fast and almost complete loss of genetic diversity in both subpopulations, which invalidates the applicability of the infinitesimal model. The process is accompanied by a further increase of the population mean of the mating trait, as selection against increased assortativity ceases. At the end of this phase, the two ecologically differentiated populations are characterized by practically complete reproductive isolation.

### Diverse sources of mismatch

Non-Gaussian shapes of the resource distribution and/or competition kernels are not the only possible sources of residual ecological selection resulting from a mismatch between resource consumption and resource production.

When the maximum allelic variance 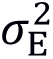 of the ecological trait is small, so is the population variance of the ecological trait. Under these conditions, the advantage of gradually increased assortativity is minimal, so the population mean of the mating trait does not increase, so the population variance of the ecological trait cannot increase either. Therefore, a strong mismatch remains between resource consumption and resource production, despite the Gaussian shapes of resource distribution and competition kernel. The population can move out of this state only through a sufficiently large fluctuation in the population mean of the mating trait. Once this happens, the second and third phase follow as before, with the three-phase speciation process being finalized by fast runaway sexual selection. These fluctuation-induced speciation dynamics are consistent with the existence of an intermediate repeller of mating-trait evolution separating two basins of attraction for the population mean of the mating trait; such repellers have been observed also in several other models (e.g., de Cara et al., 2008; Otto et al., 2008; Pennings et al., 2008). Fluctuation-induced speciation dynamics are also observed when the initial increase of the population mean of the mating trait is prevented for other reasons, e.g., for *α* = 2 (Figure S4b), which makes the consequences of an increase fourth-order small (Eq. 2c).

When the number *n*_E_ of loci for the ecological trait is small in the doubly Gaussian case, the utilization mismatch results from the fact that the distribution of resource consumption cannot be close to that of resource production, as the population distribution of the ecological trait involves only a small number of phenotype classes and is restricted to a finite phenotype interval. The study by Doebeli and Dieckmann (1999) reported deterministic speciation for *n*_E_ = *n*_M_ = 5, and these earlier results are extended in Figure 4b to significantly higher population sizes. The analyses of parameter dependences shown in Figure S7 demonstrates that, for the doubly Gaussian case, the utilization mismatch originating from a low number of loci for the ecological trait is key for deterministic, rather than fluctuation-induced, speciation dynamics.

We conclude that utilization mismatch and residual selection can arise in different ways, which can, moreover, interact with each other in complicated ways. It is beyond the scope of the present study to map all the possibilities comprehensively.

### Discretizing the continuum

At a more fundamental level, Maynard Smith and Szathmáry (1995) asked about the ultimate reason for the discreteness of species. Referring to a theoretical example of continuous coexistence reported by Roughgarden (1979, pp. 534-536), they rejected an ecological explanation of continuous coexistence and instead proposed that sexual interactions are responsible for discretizing the continuum. Noest (1997) followed this lead by deriving conditions for the instability of the sexual continuum. While our study confirms the significance of sexual selection for reinforcing reproductive isolation, we propose that the ultimate reason for the discreteness of species is ecological: a perfect match between resource production and resource consumption is non-generic.

For faithfully reproducing clonal organisms competing for a continuum of resources, the coexistence of a continuum of trait values is structurally unstable (Sasaki and Ellner, 1995; Sasaki, 1997; Gyllenberg and Meszéna, 2005; Barabás et al., 2012). While models with equilibria characterized by such continuum coexistence can easily be constructed for an arbitrary competition kernel and an arbitrary trait distribution by suitably choosing the resource distribution – with the example given by Roughgarden (1979) for Gaussian functions just being a special case –, the existence of such equilibria can always be destroyed by an arbitrarily small perturbation of the resource distribution (Sasaki, 1997). Moreover, the dynamical stability of such equilibria for uniform resource distributions requires the considered competition kernel to be positive definite (Pigolotti et al., 2007, 2010; Leimar et al., 2008; Hernández-García, et al., 2009; Sasaki and Dieckmann, 2011; Leimar et al., 2013), which can typically be destroyed by an arbitrarily small perturbation of the competition kernel. In the spirit of MacArthur (1969, 1970), we can thus say that equilibria characterized by a discrete set of trait values, instead of a continuum of trait values, represent an optimal use of resources. Except in the immediate vicinity of the non-generic case of (always) structurally unstable and (typically) dynamically unstable equilibria characterized by a continuum of trait values, the rule of thumb of ‘limiting similarity’ (MacArthur and Levin, 1967) applies: the trait difference between stably coexisting populations is approximately determined by the width of the competition kernel (Roughgarden 1974; Meszéna et al., 2006; Szabó and Meszéna, 2006; Barabás and Meszéna, 2009).

For sexually reproducing populations with multilocus genetics competing for a continuum of resources, the non-genericity of a perfect match between resource production and resource consumption under sexual inheritance is a corollary of the non-genericity of continuum coexistence under clonal inheritance. This is because the perfect uniformity of the ecological fitness landscape in a sexual population is formally equivalent to the existence of an equilibrium characterized by the coexistence of a continuum of trait values in a clonal population. Here we have demonstrated that residual selection, emerging from the generically existing utilization mismatch, can initiate adaptive speciation, resulting in ecologically differentiated species with practically complete reproductive isolation. Obviously, this transition is contingent on the availability of isolation mechanisms and on many details affecting genetics, mating, and evolution. When adaptive speciation happens, it can be considered as a manifestation of the discretization tendency inherent in ecological interactions.

### Outlook

Motivated by both empirical and theoretical studies, the growing recognition of the ecological dimensions of speciation can be seen as renewed appreciation for the original understanding of speciation proposed by Darwin (Provine, 2004; Mallet, 2008). While Darwin (1859) envisaged a gradual transformation of within-species varieties into different species driven by ecological selection (Reznick and Ricklefs, 2009), Mayr’s (1942) emphasis on the genetic dimensions of speciation had become accepted as scientific consensus for decades. While the biological species concept established a firm distinction between varieties and species, and the theory of allopatric speciation stated that no divergent evolution was possible in the presence of substantial and prolonged gene flow (see Gavrilets, 2004, for the mathematical theory), recent research re-emphasizes the occupation of new ecological opportunities, i.e., new niches, as a key driver of speciation. Accordingly, speciation should be seen as being jointly determined by ecological opportunities and genetic/physiological constraints – as any other type of adaptive evolution.

On the empirical side, it has turned out that reproductive isolation is not always strictly maintained even between established species (e.g., Beltran et al., 2002; Grant et al., 2005; Mallet, 2005; Heliconius Genome Consortium, 2012). Beyond growing evidence for sympatric speciation (e.g., Schliewen et al, 1994; Via, 2001), a large body of literature has demonstrated the role of ecological selection and prolonged gene flow during processes of speciation (e.g., Rundle and Nosil, 2005; Nosil, 2008; Niemiller et al., 2008; Nosil and Feder, 2012; Shafer and Wolf, 2013; Riesch et al., 2017). Hendry et al. (2009), Huber et al. (2007), and De León et al. (2012) have provided a possible example for ongoing ecology-driven speciation in Darwin’s finches. Seehausen (2015) and his coworkers have established the ecological basis for the adaptive radiation of cichlid fish.

On the theoretical side, there is no reason to restrict the relevance of adaptive speciation to the sympatric mode. In the context of modeling the evolution of spatial niche segregation through parapatric and allopatric speciation in spatially structured populations, Meszéna et al. (1997), Mizera and Meszéna (2003), and Szilágyi and Meszéna (2009) have studied evolutionary branching in asexual populations, while Kisdi and Geritz (1999), Geritz and Kisdi (2000), Doebeli and Dieckmann (2003), Heinz et al. (2009), Payne et al. (2011), Fazalova and Dieckmann (2012), Rettelbach et al. (2013), and Sachdeva and Barton (2017) have examined adaptive speciation in sexual populations (see Pásztor et al., 2016, Chapter 10, for an integrative perspective on niche segregation). This potential generality of the theory is in line with the increasingly influential empirically motivated notion of ecological speciation (Schluter, 2000; Nosil, 2012), which shares with the notion of adaptive speciation an emphasis on ecology-driven processes, complementing an emphasis on geographical patterns distinguished by the sympatric/parapatric/allopatric modes of speciation (see Dieckmann et al., 2004, pp. 8-9 and pp. 383-387, for integrative perspectives on classifying speciation).

Here we have tried to contribute to a deeper understanding of the interplay between ecology and genetics in speciation processes partitioning an ecological continuum based on an ecological trait given by a multilocus quantitative character. We have highlighted the importance of the residual selection pressures arising from inevitable mismatches between resource production and resource consumption, often leading to three-phase transitions to reproductive isolation. For these transitions, mixed selection regimes are key: these are simultaneously characterized by aspects traditionally associated with stabilizing and disruptive selection, selecting for increased assortativity and decreased allelic variance at the same time. Even though the processes of adaptive speciation studied here are prototypical instances of ecological speciation, sexual selection naturally emerges in these processes as a strong driver and unavoidable consequence of assortative mating: whenever speciation happens in our model, it is concluded by fast runaway sexual selection, guaranteeing the practically complete reproductive isolation of the resultant species. While we have focused this study on slow transitions to reproductive isolation, with substantial and prolonged gene flow between the incipient species, our model predicts much faster transitions for other parameter combinations (see also Nosil et al., 2017). Even the slow transitions spanning a few thousand generations are fast compared with the typical lifetime of species, in line with the notion of punctuated equilibrium (Eldredge and Gould, 1972). We conclude that the emergence of ecologically differentiated and reproductively isolated species is rooted in the structure of ecological interactions and is realized through the interplay of ecological and sexual selection.

## Acknowledgements

We thank Nick Barton, Freddy Christiansen, Michael Doebeli, Varvara Fazalova, Mark Kirkpatrick, Jim Mallet, Benjámin Márkus, Patrik Nosil, Liz Pásztor, Alexey Romanenko, and Janne-Tuomas Seppänen for insightful discussions. G.M. acknowledges support by the Hungarian Scientific Research Fund (grant K81628) and the Hungarian National Research, Development and Innovation Office (grant K123796), as well as hospitality by the Evolution and Ecology Program at the International Institute for Applied Systems Analysis.

### Appendix: Model Specification

We use a slightly modified version of the model by Dieckmann and Doebeli (1999). Dictated by the goal of studying the limit of high numbers of loci, we rescale the ecological trait values and adjust the mapping from mating trait values to mating kernels. We consider a cost of choosiness as in Doebeli and Dieckmann (2003) and investigate several metrics of segregation. The full model specification is provided below.

#### Traits

Each individual is characterized by two inherited quantitative traits: the ecological trait *x*_E_ and the mating trait *x*_M_. These are determined additively by, respectively, *n*_E_ and *n*_M_ diploid diallelic loci with equivalent allelic effects.Since each diploid locus consist of a maternally and a paternally inherited haploid locus, a trait value is the sum of 2*n*_E_ or 2*n*_M_ haploid allelic effects.

As a reference case, we are interested in the limit of infinitely many loci for the ecological trait, which requires that its allelic effects become infinitesimal in this limit (infinitesimal model, Fisher, 1918; Bulmer, 1980; Barton et al., 2017; Turelli, 2017). This limit is useful only when the population variance of the unselected trait is kept constant by suitably rescaling the allelic effects with the number of loci.

For the ecological trait, we thus scale the allelic effects with the factor 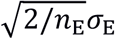,

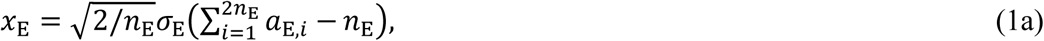

where *a*_E,*i*_ ∈ {0,1} is the allelic value at the *i*^th^ haploid locus (*i* = 1,2, …, 2*n*_E_) of the ecological trait. When the allele frequencies at all 2*n*_E_ haploid loci equal 1/2, each of these loci thus contributes 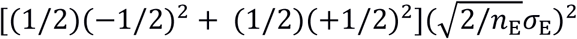 to the population variance of the ecological trait. When the allele frequencies at all 2..E haploid loci are also independent of each other, this variance equals 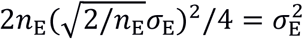, which means that 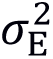 is the maximum allelic variance of the ecological trait. Note that the population variance of the ecological trait can become larger than 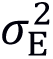 when the allele frequencies at the 2_*n*_E__ loci are positively correlated: such positive covariances contributing to the population variance of the ecological trait are a sign of linkage disequilibrium, which can be caused, e.g., by ecological selection and assortative mating. The range of the ecological trait is 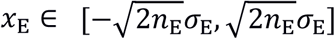, which extends to the whole real axis in the infinitesimal limit. This scaling ensures that the population variance of the ecological trait, and therefore the speed of directional evolution in the population mean of the ecological trait, as determined by Lande’s equation (Lande, 1979), remain unaffected when the number of loci for the ecological trait is varied.

For the mating trait, we scale the allelic effects with the factor 1/*n*_M_,

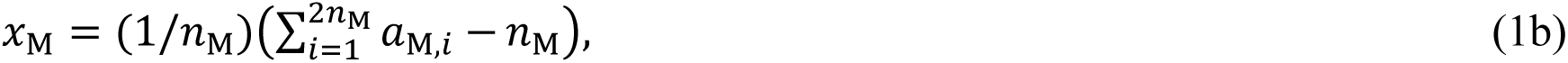

where *a*_M,*i*_ ∈ {0,1} is the allelic value at the *i*^th^ haploid locus (*i* = 1,2, …, 2*n*_M_) of the mating trait. The range of the mating trait is *x*_M_ ∈ [−1, +1], which is therefore invariant when the number *n*_M_ of loci for the mating trait is varied. The population variance of the mating trait does not play an important role in shaping the evolutionary dynamics, as the rate of directional evolution in the population mean of the mating trait is not limiting these dynamics.

#### Fecundity

Individuals are hermaphrodites. At rate *r*, an individual in its female role seeks a mate in its male role and, when successfully finding a mate, produces a single offspring. A specific individual will be chosen as mate with a probability proportional to its mating weight determined by the mating kernel *w*,

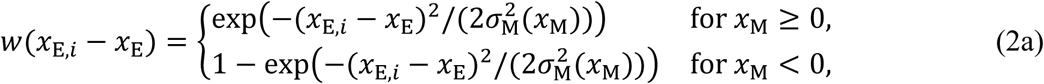

where *x*_E_ and *x*_E,*i*_, respectively, are the ecological traits of the focal individual and its potential mating partner. The standard deviation *σ*_M_ is determined by the mating trait *x*_M_ of the focal individual,

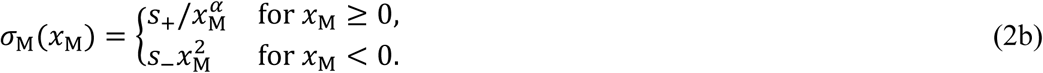

The value *x*_M_ = 0 represent random mating. Increasing positive values of *x*_M_ correspond to decreasing values of *σ*_M_ and describes strengthening assortativity, while increasing negative values of *x*_M_ describes strengthening disassortativity.

The effect of a small departure of the mating trait from *x*_M_ = 0 can be approximated as

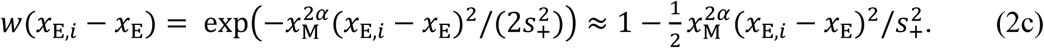

For *α* = 1/2, this effect is linear, i.e., of first order, which we therefore use for most of our study. For *α* = 2, used by Dieckmann and Doebeli (1999), this effect is quartic, i.e., of fourth order, which is unsuitable when the number of loci becomes very large.

Individuals in their female role find a mate and reproduce with probability

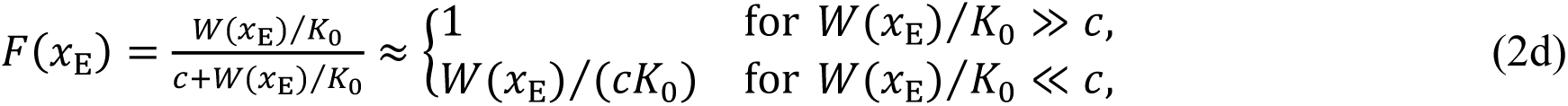

where 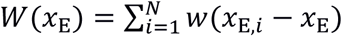 is the sum of all mating weights the focal individual with ecological trait *x*_E_ assigns to prospective mating partners, and the cost *c* of choosiness is measured by the relative total mating weight *W*(*x*_E_)/*K*_0_ for which the probability of finding a mate is halved (Doebeli and Dieckmann 2003). As the sum in *W*(*x*_E_) extends over all *N* current individuals, it is proportional to *N*; *W*(*x*_E_) is thus normalized by *K*_0_, which scales the number of individuals. Extreme choosiness results in small values of *W*(*x*_E_), entailing a low probability of reproduction. When the cost *c* of choosiness is high, the probability of finding a mate is small and proportional to *W*(*x*_E_), whereas when *c* is high, this probability saturates at 1. The value *c* = 0 corresponds to cost-free choosiness.

Free recombination is assumed between all diploid loci. At reproduction, alleles may change from 0 to 1, or vice versa, with a small mutation probability *μ* at each locus.

#### Mortality

An individual with ecological trait *x*_E_ dies at rate

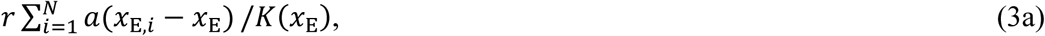

where the carrying capacity *K*(*x*_E_) at the ecological trait *x*_E_ is determined by the resource distribution *K* (see below), and the strength *a*(*x*_E,*i*_ – *x*_E_) of competition between two individuals with ecological traits *x*_E_ and *x*_E,*i*_ is determined by the competition kernel a (see below). The sum 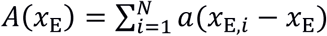 extends over all *N* current individuals and represents the competition-effective population size, in which each individual is discounted according to the strength of its competition with the focal individual with ecological trait *x*_E_.

To specify the resource distribution and the competition kernel, we use the function

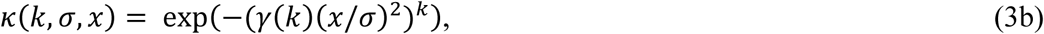

with maximum 1 at *x* = 0, standard deviation *σ*, and kurtosis parameter *k* > 0 (Figure S2; Roughgarden 1974; Pigolotti et al., 2010; Leimar et al., 2013). Here the factor

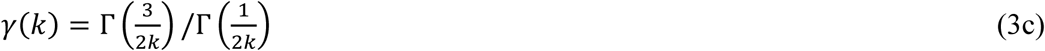

ensures that *κ*(*k, σ, x*) as a function of *x* has standard deviation *σ* for any *k* (this can be checked easily by computer algebra, e.g., by using Mathematica). The kurtosis parameter *k* = 1 yields the Gaussian function

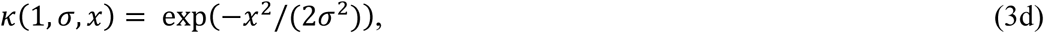

while kurtosis parameters *k* > 1 and *k* < 1 yield platykurtic and leptokurtic functions, respectively. The limiting case *k* → ∞ yields the box-shaped function

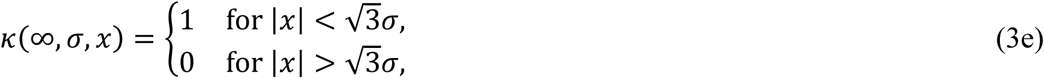

which is maximally platykurtic. The competition kernel a is given by

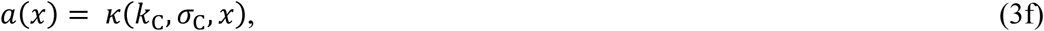

with kurtosis parameter *k*_C_ and standard deviation *σ*_C_, and the resource distribution *K* is given by

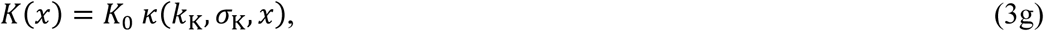

with kurtosis parameter *k*_K_, standard deviation *σ*_K_, and maximum *K*_0_; the latter scales the total number of individuals.

#### Genetic diversity

The allelic variance within the phenotype class *x*_E_ of the ecological trait is

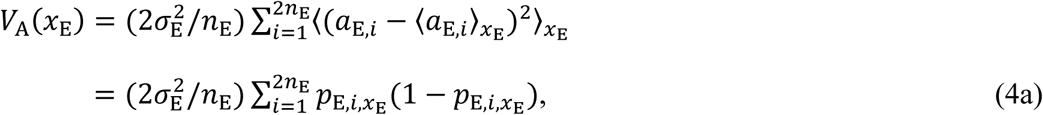

where 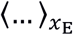 denotes the average over all individuals with ecological trait *x*_E_ and

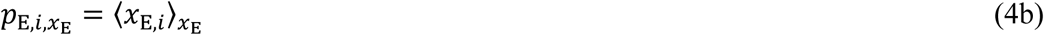

is the allele frequency within the phenotype class *x*_E_ at the haploid locus *i* of the ecological trait.

For both the ecological trait and the mating trait, the rates of directional evolution in the allele frequencies of individual loci go to zero as the allelic effects of individual loci go to zero in the limit of infinitely many loci. For a finite number of loci, the selection-driven change of allelic variance thus becomes slow when the number of loci increases. For infinitely many loci, i.e., in the infinitesimal model, the change of allelic variance becomes infinitely slow, i.e., arrested, and thus cannot be affected by selection.

#### Fitness

The fitness of individuals in phenotype class *x*_E_ of the ecological trait is determined by the difference between their birth rates and death rates,

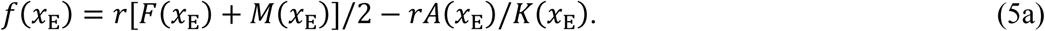

The first term above is the per capita birth rate, averaged among individuals in their female and male roles. The birth rate is proportional to *r*, as this is the rate at which females seek mates and, if successful, reproduce. For individuals in their female role, this is multiplied by the probability *F*(*x*_E_) of finding a mate, while for individuals in their male role, this is multiplied by the propensity *M*(*x*_E_) of being chosen as a mate. The factor 1/2 arises since each offspring has a mother and a father. The second term above is the per capita death rate, which is independent of whether individuals are in their female or male role.

We can partition this fitness into components associated with ecological selection and sexual selection, *f*(*x*_E_) = *f*_e_(*x*_E_) + *f*_s_(*x*_E_), with the ecological fitness

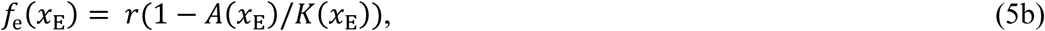

the sexual fitness

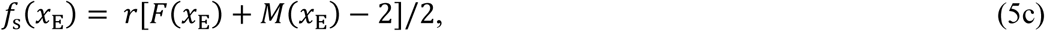

and the male reproductive success

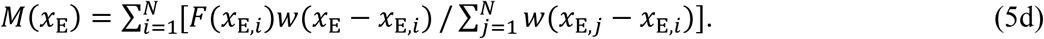

When choosiness is cost-free, *F*(*x*_E_) = 1 and sexual fitness is a linear function of male reproductive success.

#### Segregation indicators

To follow salient changes in the population’s mating dynamics throughout three-phase transitions to reproductive isolation, we use several indicators of segregation: the average squared difference between the ecological traits of parents, the average number of loci at which the ecological genotypes of the two parents differ (average Hamming distance), and the average squared difference of an offspring’s ecological trait from the mid-parental ecological trait of its parents (segregation variance).

### Supplementary Figures

**Figure S1.**
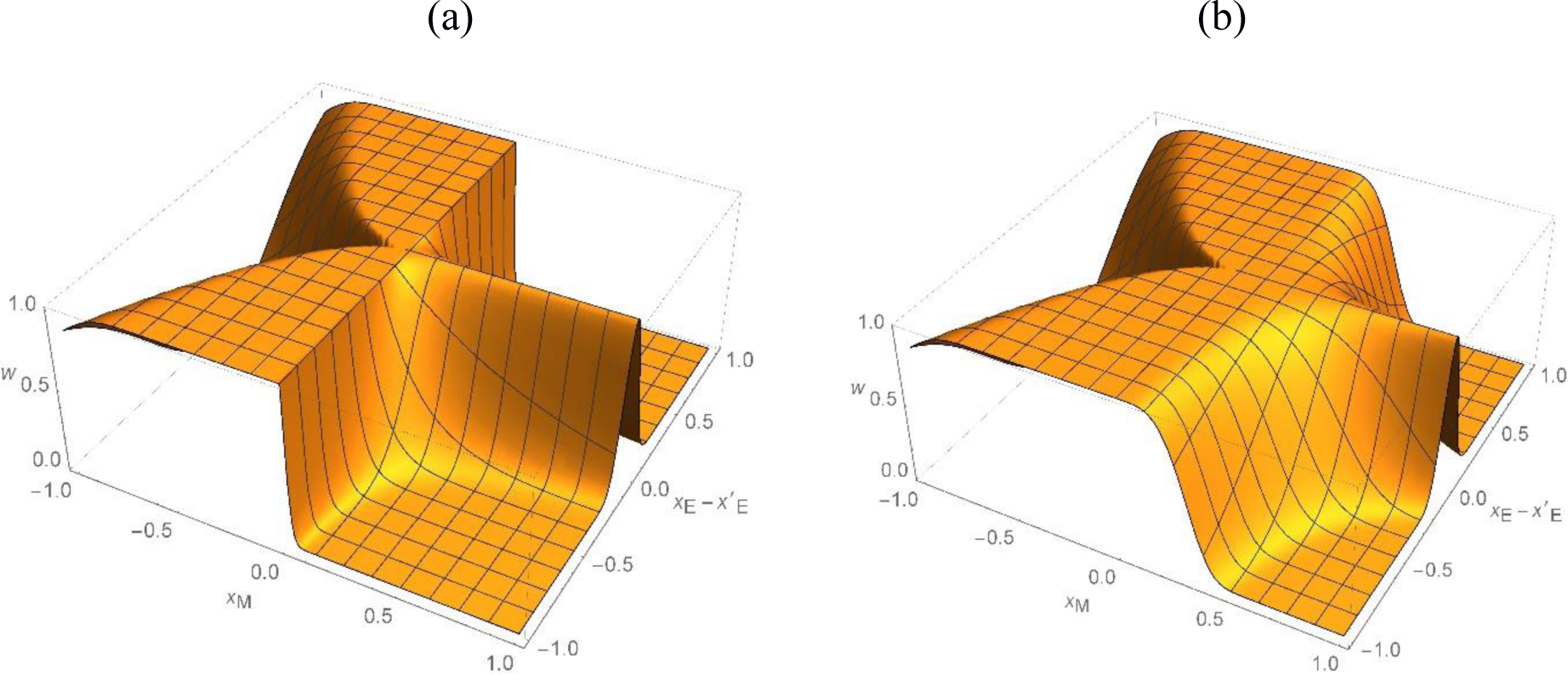
Shapes of the function *w* (Eqs. 2a and 2b) used to describe mating weights in dependence on the female mating trait *x*_M_ and the difference between the male and female ecological traits *x*_E_, for (a) *α* = 0.5 and (b) *α* = 2.

**Figure S2.**
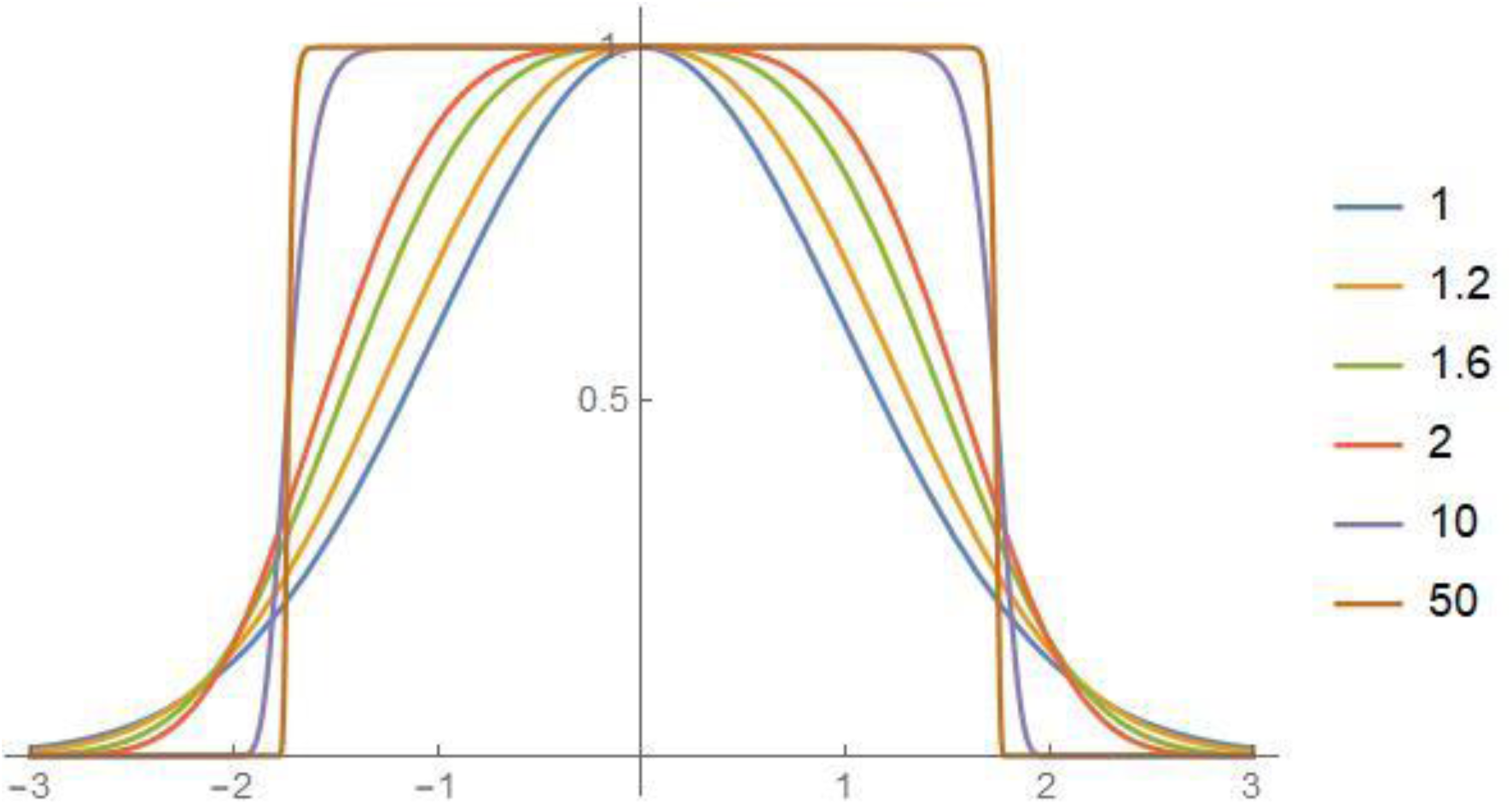
Shapes of the function *κ* (Eq. 3b) used to describe the resource distribution and the competition kernel, for different values of its kurtosis parameter *k*. This function is Gaussian for *k* = 1 and becomes more and more box-shaped as *k* is increased.

**Figure S3.**
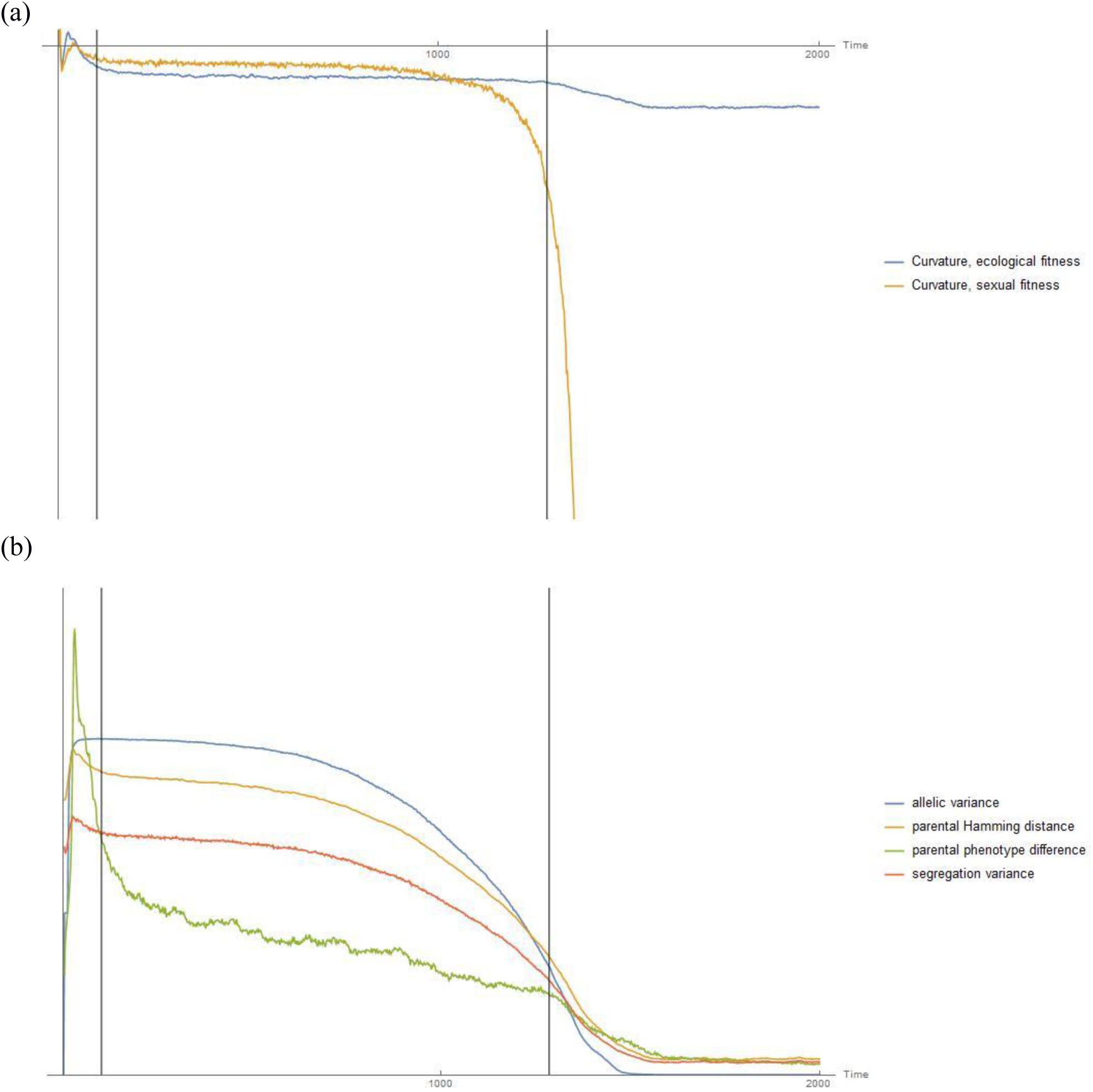
Time series of summary statistics describing the course of the three-phase transition to reproductive isolation in Figure 1. The boundaries between the three phases are indicated by vertical lines. (a) Curvatures describing the fitness landscape: average curvature of the ecological fitness (blue) and average curvature of the male reproductive success (brown), at correct relative scale. (b) Segregation indicators describing mating pairs: allelic variance of the ecological trait averaged over the two phenotype classes with maximum abundance at the end of the speciation process (blue), average Hamming distance between the genotypes of the ecological traits of mating pairs (brown), average squared difference between the ecological traits of mating pairs (green), and segregation variance (red), at arbitrary relative scales.

**Figure S4.**
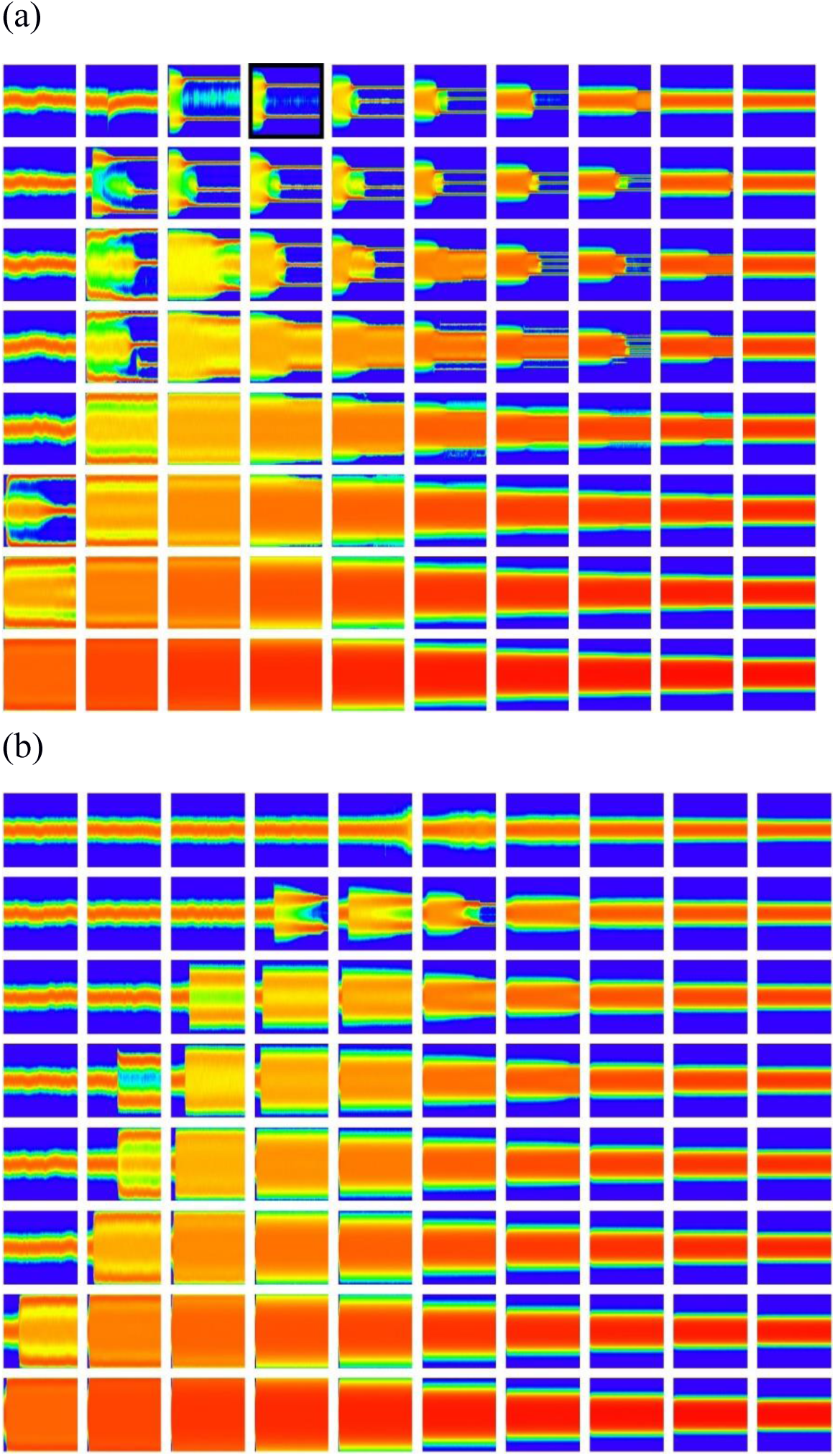
Dependence of three-phase transitions to reproductive isolation on the standard deviation *σ*_C_ of the competition kernel, the variance parameter *σ*_E_ of the ecological trait, and the parameter *α*. In each panel, the horizontal coordinate is time, the vertical coordinate is the ecological trait, and the color coding indicates the population distribution of the ecological trait. In each of the two matrices of panels, *σ*_C_ = 0.1, 0.2, 0.3, 0.4, 0.5, 0.6, 0.8, 1 increases in the rows of panels from bottom to top, while *σ*_E_ = 0.1, 0.15, 0.2, 0.25, 0.3, 0.4, 0.5, 0.6, 0.7, 0.8 increases in the columns of panels from left to right. In (a) *α* = 1/2, while in (b) *α* = 2. Other parameters are as in Figure 1, except for *K*0 = 10^4^ and *T* = 5,000. The panel framed in black indicates the combination of parameters *σ*_C_, *σ*_E_, and *α* used in Figure 1.

**Figure S5.**
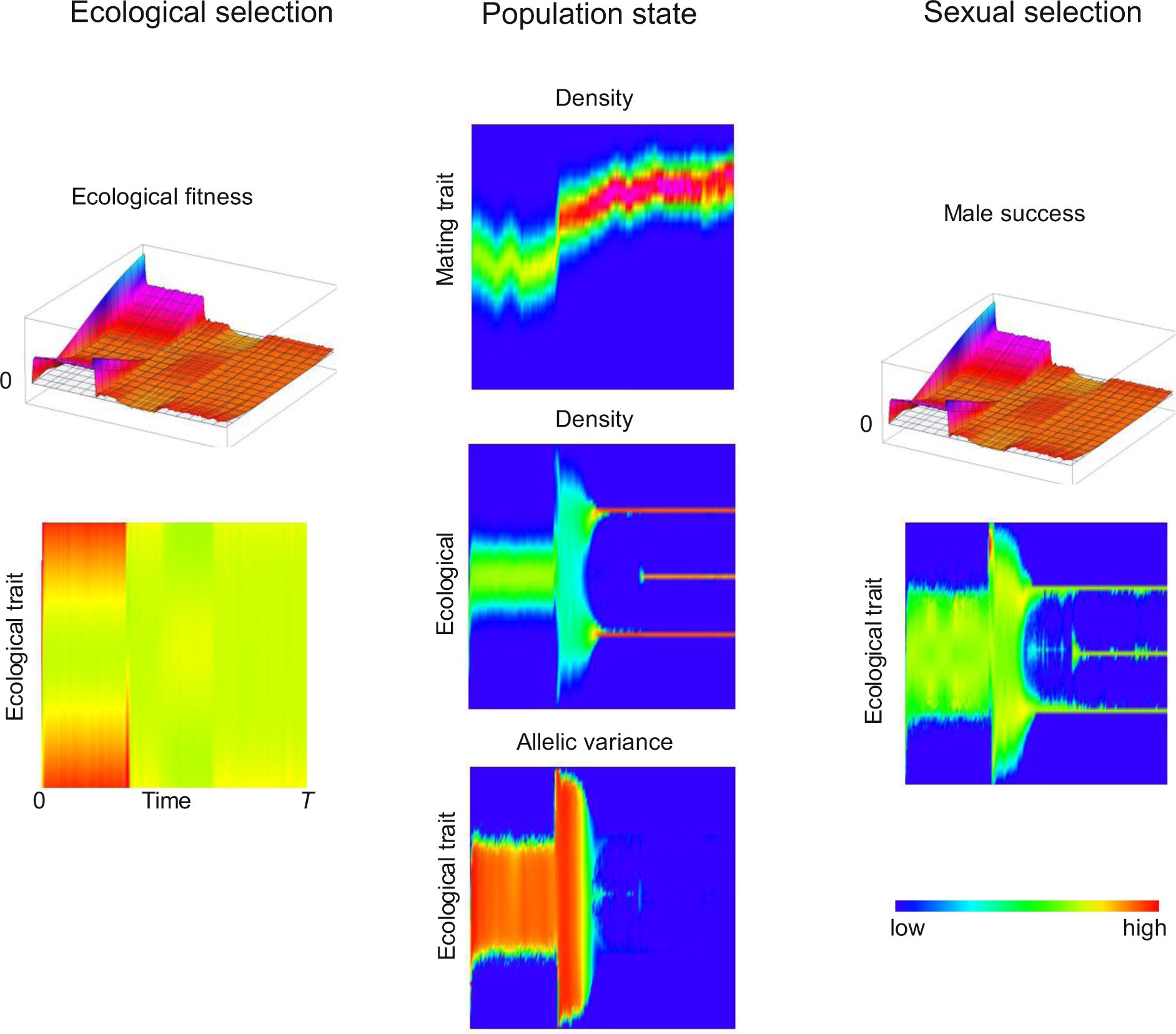
Three-phase transition to reproductive isolation in the doubly Gaussian case in Figure 4a, shown at the same level of detail as in Figure 1. The second phase is initiated by a fluctuation-induced transition to high population variance in the ecological trait. The degree of demographic stochasticity in this figure is higher than in Figure 1 because of the ten-fold smaller population size.

**Figure S6.**
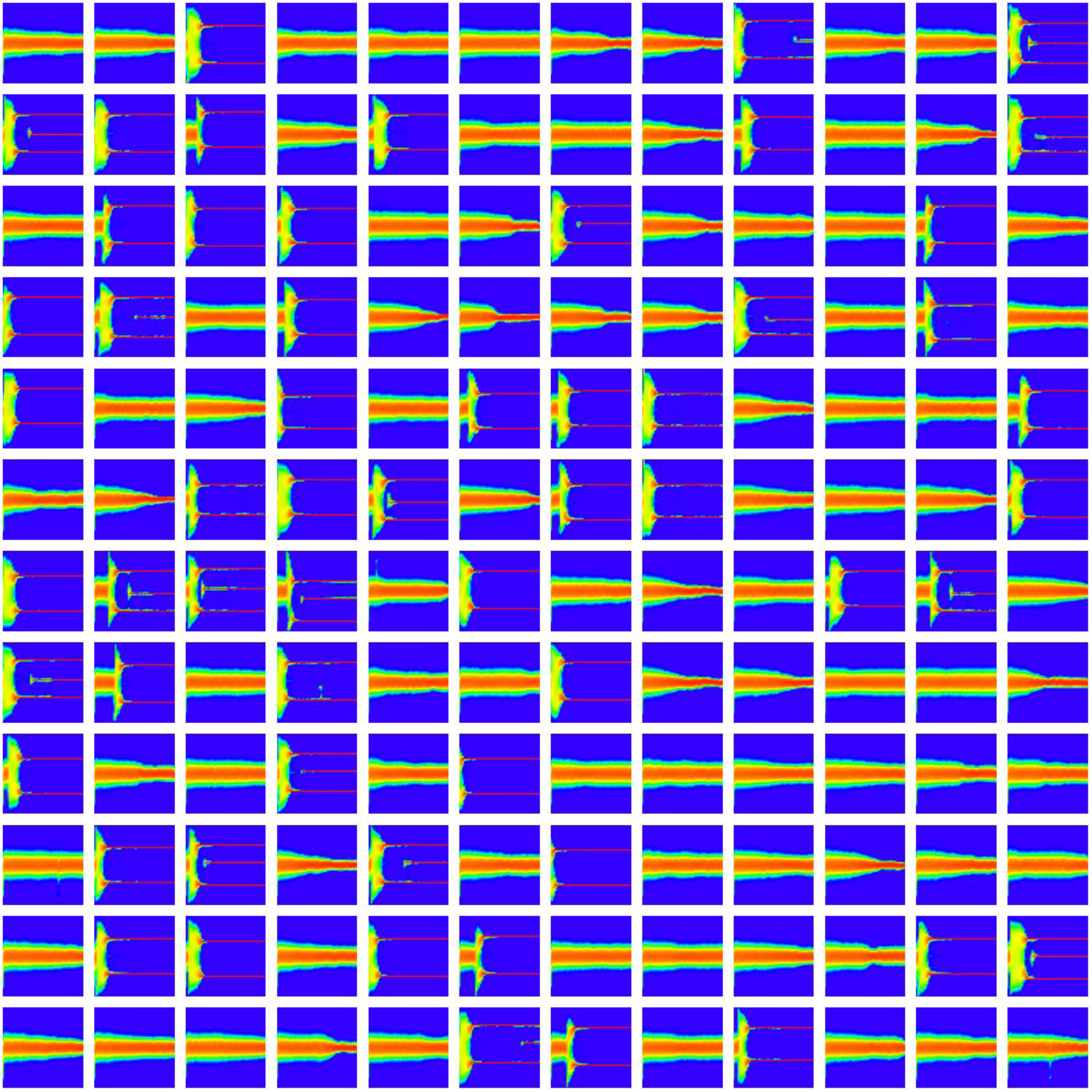
Replicate model runs for the doubly Gaussian case with the same parameter values as in Figures 4a and S5. The 144 shown realizations thus differ only in their random seed. As the population tends to lose allelic variance in the ecological trait while it stays unimodal, the fluctuation-induced transition to high population variance in the ecological trait has a finite time window to occur.

**Figure S7.**
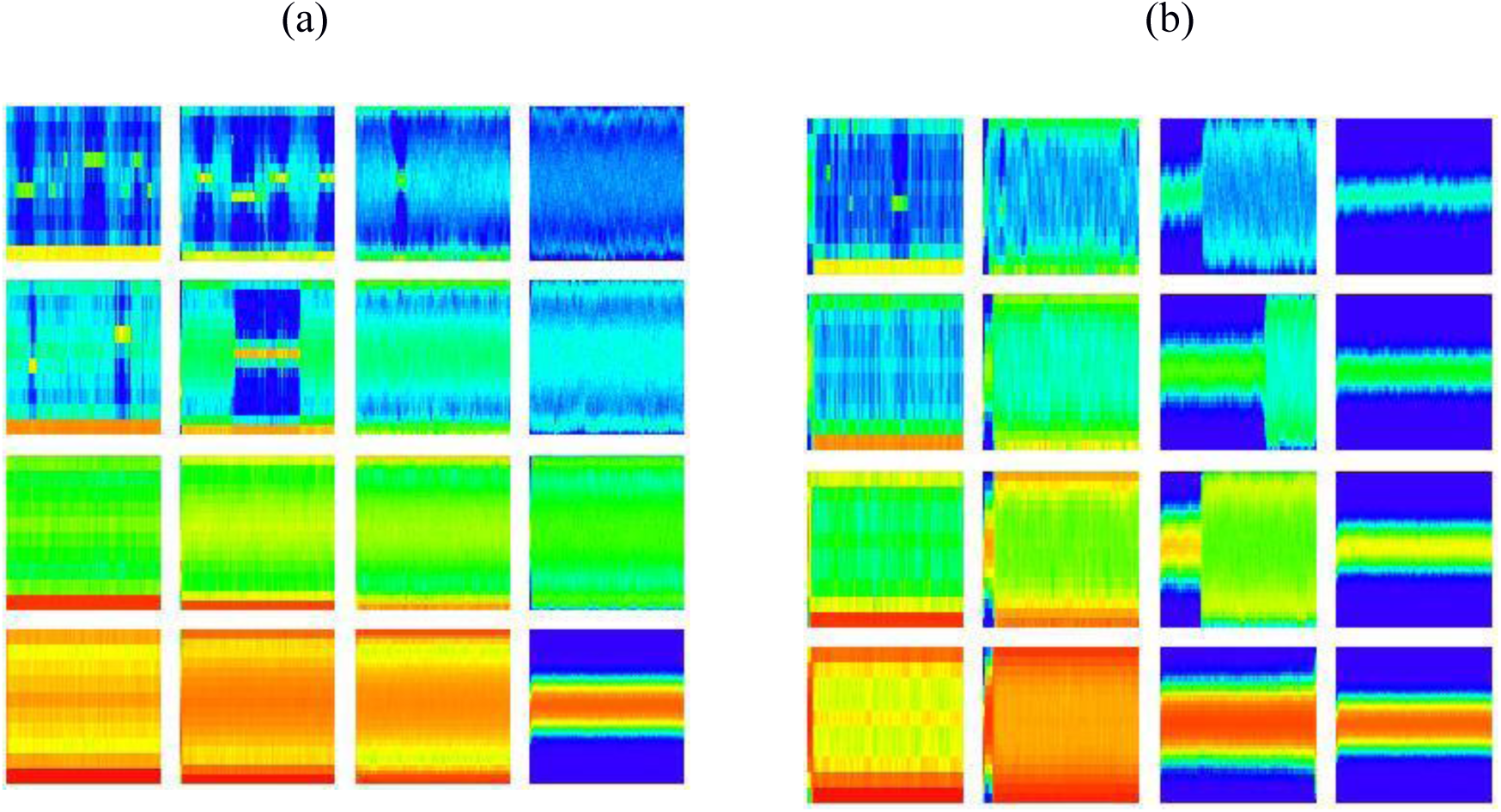
Dependence of the dynamics in the doubly Gaussian case on population size, the numbers of loci, and the parameter *α*. In each panel, the horizontal coordinate is time, the vertical coordinate is the ecological trait, and the color coding indicates the population distribution of the ecological trait. In each of the two matrices of panels, *K*_0_ = 500, 1,000, 3,000, 10,000 increases in the rows of panels from top to bottom, while *n*_E_ = *n*_M_ = 5, 8, 16, 32 increase in the columns of panels from left to right. In (a) *α* = 1/2, while in (b) *α* = 2. Other parameters are as in Figure 1, except for *μ* = 10^−3^ and *T* = 5,000.

